# A Spatiotemporal Compartmentalization of Glucose Metabolism Guides Mammalian Gastrulation Progression

**DOI:** 10.1101/2023.06.06.543780

**Authors:** Dominica Cao, Liangwen Zhong, Anupama Hemalatha, Jenna Bergmann, Andy L. Cox, Valentina Greco, Berna Sozen

## Abstract

Gastrulation is considered the *sine qua non* of embryogenesis, establishing a multidimensional structure and the spatial coordinates upon which all later developmental events transpire. At this time, the embryo adopts a heavy reliance on glucose metabolism to support rapidly accelerating changes in morphology, proliferation, and differentiation. However, it is currently unknown how this conserved metabolic shift maps onto the three-dimensional landscape of the growing embryo and whether it is spatially linked to the orchestrated cellular and molecular processes necessary for gastrulation. Here we identify that glucose is utilised during mouse gastrulation via distinct metabolic pathways to instruct local and global embryonic morphogenesis, in a cell type and stage-specific manner. Through detailed mechanistic studies and quantitative live imaging of mouse embryos, in parallel with tractable *in vitro* stem cell differentiation models and embryo-derived tissue explants, we discover that cell fate acquisition and the epithelial-to-mesenchymal transition (EMT) relies on the Hexosamine Biosynthetic Pathway (HBP) branch of glucose metabolism, while newly-formed mesoderm requires glycolysis for correct migration and lateral expansion. This regional and tissue-specific difference in glucose metabolism is coordinated with Fibroblast Growth Factor (FGF) activity, demonstrating that reciprocal crosstalk between metabolism and growth factor signalling is a prerequisite for gastrulation progression. We expect these studies to provide important insights into the function of metabolism in other developmental contexts and may help uncover mechanisms that underpin embryonic lethality, cancer, and congenital disease.

## Introduction

The events that shape the creation of a differentiated multicellular structure from a single totipotent zygote all coalesce during a fundamental developmental transition known as gastrulation. At this stage, the three germ layers (ectoderm, mesoderm, endoderm) are specified and spatially organized along key axes to form the primordia of major organs and tissues (Arnold and Robertson, 2009). Gastrulation is initiated by an evolutionarily conserved set of cell movements and signalling pathways at the embryo’s posterior pole and gives rise to the primitive streak (PS) — a dynamic structure that forms all mesoderm tissues (Rivera-Perez and Hadjantonakis, 2014). During this process, embryonic epiblast cells exit pluripotency and undergo a sequence of transcriptional and morphological changes that result in an epithelial-to-mesenchymal transition (EMT) to ingress into the PS (Tam and Beddington, 1992) **(Figure 1A)**. Nascent mesoderm cells specified within the PS then migrate laterally around the embryonic circumference in an organised fashion until they reach the anterior pole, transforming the embryo from a cellular mass into a multi-layered organism **(Figure 1A)**. Localised sources of morphogen signals promote diverse functions, orchestrating cell fate decisions and morphogenetic behaviours to shape the embryo (Bardot and Hadjantonakis, 2020). How these critical signals integrate at the correct time and place to mediate gastrulation morphogenesis remains a long-term goal in the field of developmental biology, and the remarkable fidelity of this process suggests that multiple layers of regulation are required to ensure robust embryonic patterning.

**Figure 1.**
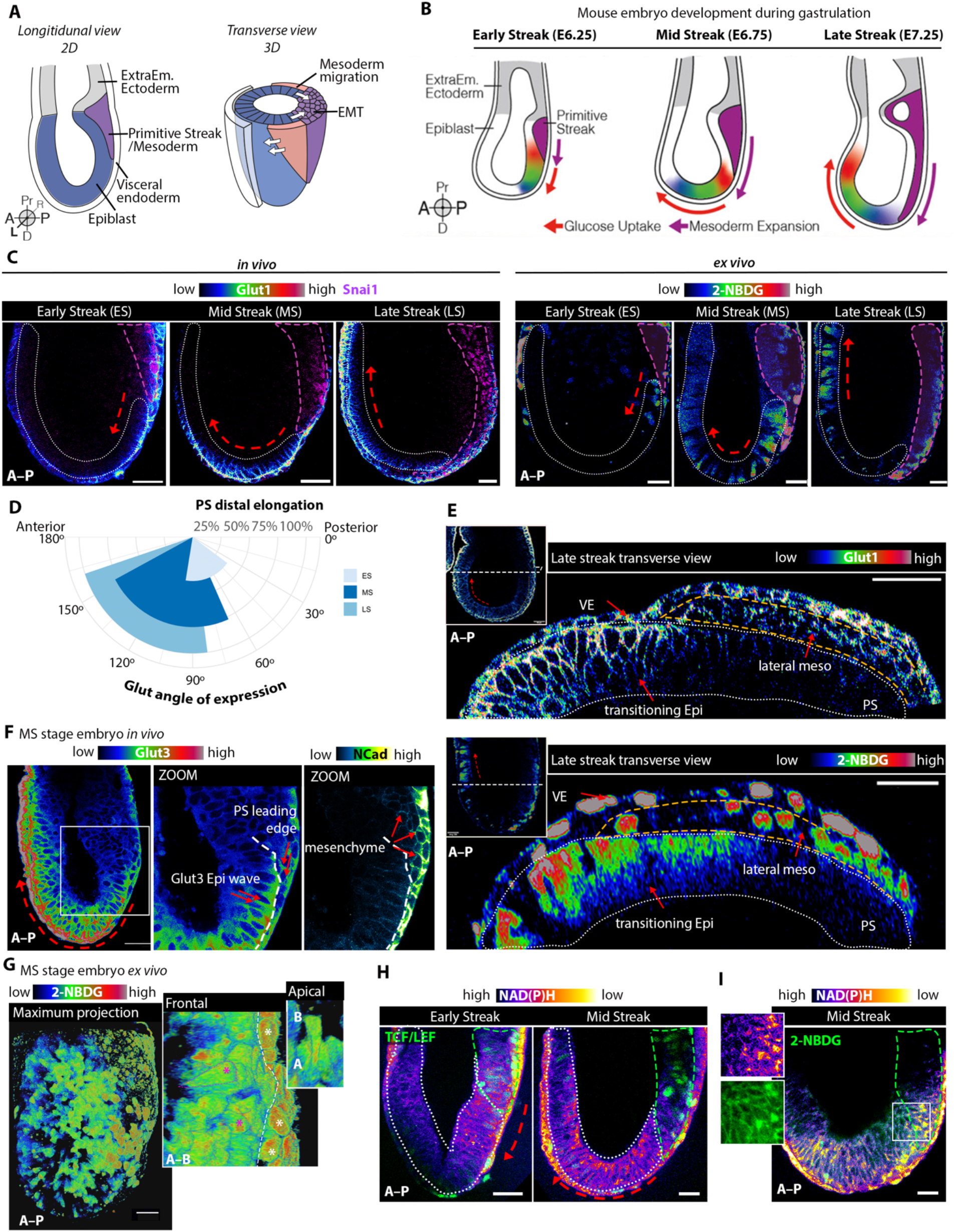
A wave of glycolytic activity precedes mouse gastrulation. **(A)** Schematic of the gastrulation stage mouse embryo. **(B)** Schematic of the glucose-uptake ‘wave’ phenotype throughout the progressive stages of mouse gastrulation. **(C)** Single Z-sections of: **Left:** *in utero*-dissected gastrula stage embryos stained for Glut1 and Snai1 (n=55 embryos: ES n=8, MS n=26, LS n=19); **Right:** gastrula stage embryos following 2h of *ex-vivo* incubation with 2-NBDG (n=27 embryos: ES n=5, MS n=13, LS n=9). Epiblast, dotted-white regions; PS, dotted-magenta regions. Scale bars represent 40µm. Heatmap intensity colours used via Fiji’s LUT. **(D)** Fig.1B-C quantifications of the mean angle of Glut1/3 expression in the epiblast, plotted against the mean of the PS’s distal length. As development progresses, the region of epiblast Glut expression extends anteriorly. **(E)** Orthogonal transverse sections showing a wave of Glut1 and 2-NBDG expression in epiblast (dotted-white regions) and lateral mesodermal wings (dotted-yellow regions). Scale bars represent 40µm. Heatmap intensity colours used via Fiji’s LUT. **(F)** *In utero*-dissected MS embryo stained for Glut3 and Ncad (n=15 embryos). Scale bars represent 40µm. Heatmap intensity colours used via Fiji’s LUT. **(G)** 3D-reconstructed images of a 2-NBDG incubated MS embryo, revealing a gradient of glucose uptake in epiblast (pink asterisks) and migratory mesoderm (white asterisks). Heatmap intensity colors used via Fiji’s LUT. **(H)** Live multi-photon imaging of TCF/LEF-GFP (marks primitive streak) reporter embryos for endogenous NAD(P)H, as a readout of glycolytic activity (n=12 embryos: ES n=7, MS n=5). Epiblast, white-dotted regions; and PS, green-dotted regions. Scale bars represent 40µm. **(I)** Live multi-photon NAD(P)H imaging of embryos following 2h incubation with 2-NBDG (n=2 MS embryos). Scale bars represent 40µm.

To explain how an organism can establish positional identity during development and regeneration, a “gradient theory” – whereby graded metabolism along an embryonic axis directs tissue patterning and morphogenesis – was first experimentally introduced in 1915 (Child, 1941). While this model had been challenging to incorporate into a mechanistic framework, it is now supported by the recent conceptualization of *metabolic signalling* – where metabolic enzymes function beyond bioenergetics to actively modulate or instruct cellular and developmental programs (Miyazawa and Aulehla, 2018). Indeed, rising interest across multiple fields has highlighted the degree to which metabolic pathways can act as developmental regulators. For example, regionalized glycolytic gradients have been described for both chick and mouse in the developing tail bud during mid-gestation (Bulusu et al., 2017; Oginuma et al., 2017), and evidence also indicates that glycolysis can function independently of energy production during epiblast-derived tissue development, as shown in retina (Agathocleous et al., 2012), neural crest (Bhattacharya et al., 2020), and during embryonic stem cell differentiation (Carey et al., 2015; Cliff et al., 2017; Moussaieff et al., 2015). Despite its critical importance, it is still unclear how metabolic signalling might be linked to morphogen signalling pathways, and how it might modulate cell function more generally is poorly understood and remains unexplored in the context of early post-implantation morphogenesis.

A lack of accessible methods to simultaneously probe metabolic and morphogen signalling kinetics at the single cell scale within the context of live, gastrulating embryos has hindered efforts to understand how these pathways might communicate to support complex differentiation and morphogenetic events. Here, we address these challenges through combined *in vivo* and *ex vivo* live imaging of gastrulating mouse embryos, as well as with support from *in vitro* stem cell and embryo-derived tissue explant models. Using these tools, we dissect the instructive role of cellular metabolic activity in time and space for the coordination of mouse gastrulation. We define a spatiotemporally-regulated pattern of glucose uptake that arises initially in the mouse early post-implantation epiblast and reappears in lateral mesenchymal cells, and find that this phenotype serves a functional role in supporting three key fundamental aspects of gastrulation: pluripotency exit, EMT, and mesoderm migration. Importantly, our data show that these processes rely on distinct glucose metabolizing pathways. We show that while the HBP is specifically crucial to support cell-state transitions, glycolysis becomes necessary during mesenchymal migration. Furthermore, we find that these glucose-mediated events coincide with high FGF activity and suggest that an intimate relationship exists between glucose metabolism and FGF signalling to drive successful tissue patterning during mammalian gastrulation. Collectively, our integrated examination of metabolic and signalling axes provides a comprehensive framework for understanding the mechanisms that guide gastrulation, one of the most fundamental and evolutionarily conserved stages of animal life.

## Results

### A wave of glucose uptake guides mouse gastrulation onset and progression

It is well established that a major metabolic transition occurs around the time of embryo implantation, where epiblast cells switch from bivalent respiration (glycolysis and oxidative phosphorylation) towards a primarily glycolytic programme (Malkowska et al., 2022). However, it is currently unknown how this metabolic switch maps onto the changing landscape of post-implantation mammalian embryogenesis. To explore how glucose metabolism is regulated over the course of gastrulation, we first analysed in utero dissected embryos for expression of the canonical glucose transporter, Glut1, and performed a live glucose uptake analysis using fluorescent glucose (2-NBDG, (Zou et al., 2005)) in ex vivo cultured embryos as they develop through early, mid and late streak stages of gastrulation (E6.25-6.5, ES; E6.5-6.75, MS; E6.5-7.0, LS) **(Figure 1B)**. Both Glut1 protein expression and the ratiometric expression of 2-NDBG uptake demonstrated a striking antero-posterior gradient, revealing a wave of glucose uptake within the developing epiblast **(Figures 1B, C, D)**. During gastrulation onset and formation of the primitive streak (PS) in ES stage gastrulas, high glucose uptake was observed within epiblast cells at the posterior-proximal-most embryonic pole **(Figures 1B, C, D)**. As gastrulation proceeds and the PS elongates distally, this pattern of glycolytic activity extends towards the anterior-distal axis within epiblast tissue, whereas cells within the PS show no or minimum glucose uptake. Simultaneously, Glut1 expression becomes gradually reduced as cells enter the streak **(Figures 1C-E, S1A-C)**. This was also observed in the expression pattern of another canonical glucose transporter, Glut3 **(Figures 1F, S1D)**. Interestingly, we found that cells switch back to a glycolytic programme after exiting the PS as migratory mesenchyme, with high glucose metabolic activity observed within the gastrula’s lateral mesodermal ‘wings’ **(Figures 1E-G)**. Of note, although the visceral endoderm (VE) layer showed high glycolytic activity, activity levels were global across all VE cells and not spatiotemporally-resolved as observed in the epiblast **(Figure 1E)**.

To further validate these findings and expand our phenotypic characterizations, we employed a live imaging approach that utilizes multiphoton microscopy to visualize NAD(P)H, an endogenous auto-fluorescent readout of glycolytic activity (see methods) (Hemalatha et al., 2022; Quinn et al., 2013; Skala et al., 2007), in developing mouse gastrulas from a H2B-GFP:Tcf-LEF reporter mouse line (Ferrer-Vaquer et al., 2010). In support of our earlier findings, we find that NAD(P)H is also intrinsically graded over the course of gastrulation and localized to epiblast cells anterior to the expanding H2B-GFP:Tcf-LEF-expressing PS population of mesendoderm progenitors **(Figure 1H)**. Further, NAD(P)H intensity gradient overlapped with regions of 2-NBDG uptake, confirming this assay’s specificity in imaging the metabolic activity of live gastrulas (**Figure 1I)**.

Finally, we probed a spatial transcriptome dataset of the mouse gastrula (Peng et al., 2016) for key genes involved in the glycolytic pathway. The resulting gene expressions were regionalised to epiblast, and included *Slc2a1, Gpi1, Pfkb*, and *Ldhb* during progressive stages of gastrulation. Notably, the expression patterns of these genes showed similar spatial trends in relation to the expression of the PS marker *T* **(Figure S1E)**. Collectively, these data highlight a spatially and temporally resolved wave of glucose uptake that initiates in the posteriorly positioned PS at the onset of mouse gastrulation. This metabolic activity extends anteriorly in epiblast tissue as development progresses, and re-emerges within the mesenchymal cells that exit the streak and migrate anteriorly within the mesodermal wings.

### Glucose metabolism via HBP is necessary for mesoderm fate acquisition and maintenance

To systematically determine the importance of glucose metabolism for gastrulation progression and patterning, we treated ex vivo developing mouse embryos with specific chemical inhibitors that block different enzymatic steps of glucose metabolism (**Figure 2A**). For these experiments, we first used 2-deoxy-D-glucose (2-DG) and 3-bromopyruvate (BrPA), competitive inhibitors of the hexokinase (HK) and glucose phosphate isomerase enzymes (Barban and Schulze, 1961) (**Figure 2A**). ES stage gastrulas treated with 2-DG + BrPA for 18h were significantly delayed in their developmental progression. The majority of embryos failed to progress past the LS stage, while controls were able to successfully develop through post-gastrulation stages **(Figures 2B-C)**. As we observed that 18h cultured control embryos were successful in forming double mesoderm wings, we then performed shorter treatments, confined to 12h, in order to define earlier effects on PS progression. As such, this revealed that glucose metabolism inhibition impairs distal elongation of the PS **(Figures 2D, E)**.

**Figure 2.**
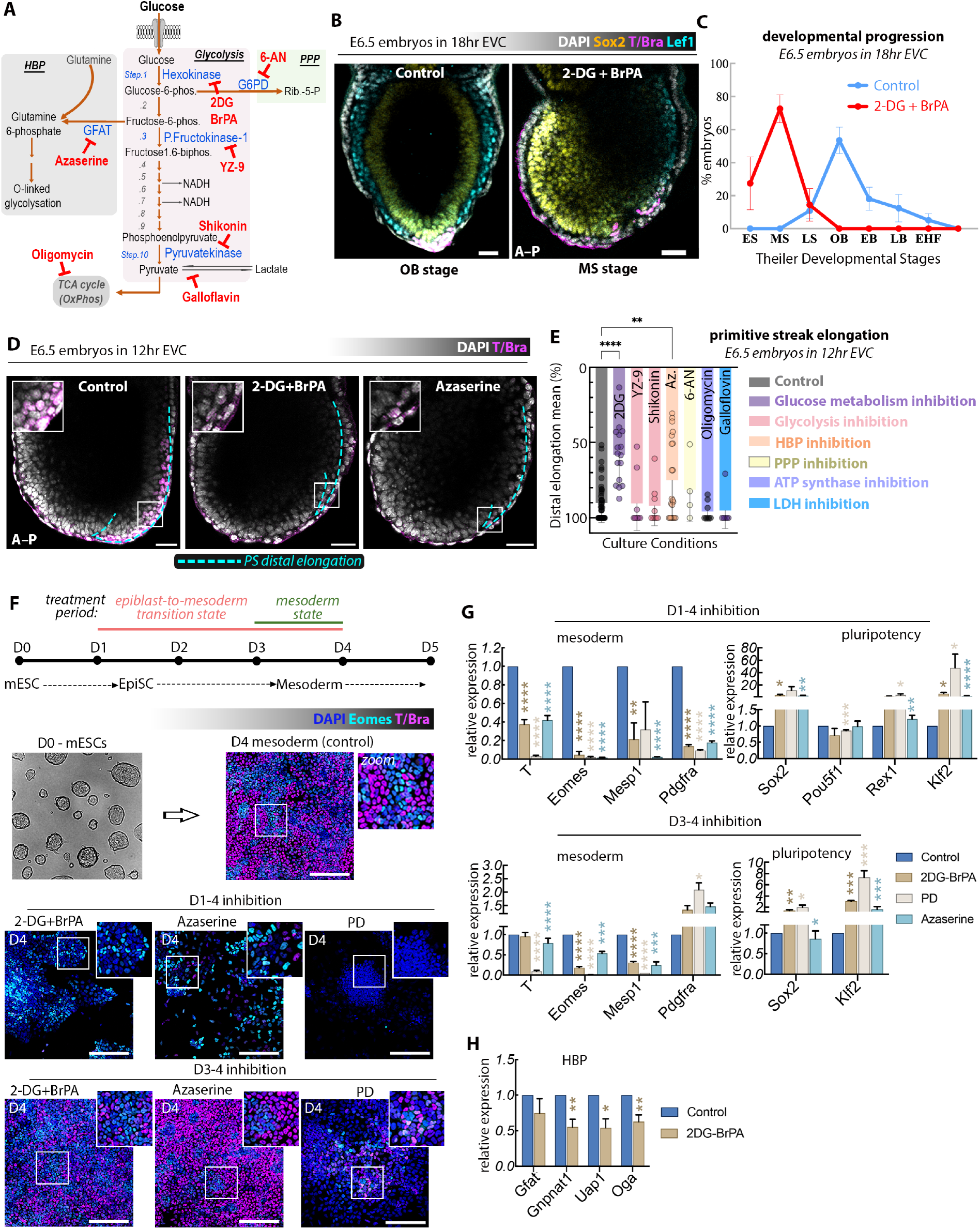
Glucose metabolism via HBP is necessary for mesoderm fate acquisition and maintenance. **(A)** Schematic of glucose metabolism and its three branches: grey = Hexosamine Biosynthetic Pathway (HBP); pink = core/late-stage glycolysis; green = Pentose Phosphate Pathway (PPP). Red text indicates chemical inhibitors and their metabolic targets in blue. **(B)** Representative Z-sections showing the expression pattern of Sox2, T/Bra, and Lef1 in embryos treated with 2-DG + BrPA for 18h *ex vivo* and **(C)** their Theiler developmental outcomes: Control (n=17), 2-DG + BrPA (n=24) across 6 experimental replicates; early streak (ES), mid-streak (MS), late streak (LS), no bud (OB), early bud (EB), late-bud (LB), early head fold (EHF). Scale bars represent 40µm. Plots show mean ± SEM. On average, ∼53% of control embryos develop to OB stage gastrulation, while ∼73% of 2-DG + BrPA treated embryos only develop to MS stage gastrulation. **(D)** Representative Z-sections of 2-DG + BrPA and 5µM Azaserine-treated embryos following 12h *ex vivo* culture (EVC). Inhibiting glycolysis impairs PS elongation (cyan dotted-lines) and reduces T/Bra expression (cyan). Scale bars represent 40µm. **(E)** Comparison of PS distal elongation lengths in embryos treated with various metabolic inhibitors: Control (n=44), 2-DG + BrPA (n=16), YZ9 (n=9), 5µM Shikonin (n=9), 5µM Azaserine (n=21), 2µM 6-AN (n=4), 10µM Oligomycin (n=8), Galloflavin (n=6). Plots show mean ± SEM. Two-tailed parametric t-test. **P < 0.01, ****P < 0.0001. **(F) Top:** Schematic of mesoderm directed-differentiation steps from mESCs. **Bottom:** 2-DG + BrPA, Azaserine, and PD-treated groups show a significant absence of Eomes (cyan) and T/Bra (magenta). Scale bars represent 100µm. 3 independent experimental replicates. **(G)** qPCR analyses of directed-differentiation experiments, querying transcript changes in mesodermal, pluripotency (treatment from day1-4 or day3-4), and **(H)** HBP genes (treatment from day1-4) across 3 independent experimental replicates. Plots show mean ± SEM. Two-tailed parametric t-test. *P < 0.05, **P < 0.01, ****P < 0.0001.

Notably, this phenotype was recapitulated with Azaserine treatments, which inhibits the HBP branch of glycolysis (Marshall et al., 1991) **(Figures 2A, D, E)**, HBP is a nutrient-sensing pathway that produces the UDP-GlcNac amino sugar, making this finding particularly interesting, as less than 1% of glucose-derived carbons are expected to be utilized through the HBP branch for the production of UDP-GlcNac (Olson et al., 2020). In contrast, blocking glycolytic enzymes that bypass the HBP branch – which we refer to as ‘late-stage-glycolysis’ components – with YZ9 (targeting PFKFB3 (Chi et al., 2020)) or Shikonin (targeting pyruvate kinase M2 (Chen et al., 2011)) had minimal or no impact on distal PS elongation **(Figures 2A, E, S2A)**. Additional perturbations using galloflavin (which targets lactate dehydrogenase (LDH) (Manerba et al., 2012)) and 6-AN (which targets PGD of the pentose phosphate pathway (PPP) (Kohler et al., 1970)) also had no significant effect on streak elongation **(Figures 2E, S2A)**. Furthermore, oligomycin-based inhibition of ATP synthase, which is necessary for oxidative phosphorylation of ADP to ATP (Cliff et al., 2017), also yielded no significant elongation defect **(Figures 2E, S2A)**.

As the highly conserved HBP branch generates substrates important for protein post-translational modifications, such as glycosylation and GlcNAcylation (Czajewski and van Aalten, 2023; Tian et al., 2012), we reasoned that glucose metabolism during early primitive streak induction might have critical roles beyond bioenergetics, in particular as a critical component for signal transduction. Importantly, neither 2-DG + BrPA nor Azaserine treatments impeded cell proliferation within developing embryos, further supporting our hypothesis that HBP may play a significant role in linking nutrient availability to gene regulation and cell signalling during early mouse gastrulation **(Figure S2B)**.

We next sought to functionally validate these ideas by culturing mouse gastrulas in nutrition-sparse media free of glucose, pyruvate, and glutamine. Expectedly, these embryos were significantly delayed in their development and exhibited impaired distal PS elongation, with no loss of Sox2 expression within the epiblast **(Figure S2C)**. Upon selective reintroduction of the excluded nutrients, we found that the addition of glucose and/or glutamine – both of which feed into the HBP wing of glycolysis– was able to fully rescue the distal PS elongation of these developing gastrulas **(Figure S2C)**.

To determine the specific role of glucose metabolism, especially HBP, for early mesoderm development and progression, we performed mesoderm-directed differentiation (Gouti et al., 2014) of mouse embryonic stem cells (mESCs) in vitro and analysed whether metabolic inhibition hinders pluripotency exit and/or mesodermal-fate acquisition and maintenance. We applied selective inhibitor treatments from day 1 to day 4 to target the epiblast-to-mesoderm transition state, or from day 3 to day 4 to target cells that had already reached the mesodermal state (**Figure 2F)**. Similar to our earlier findings with bona fide developing embryos, we found that 2-DG + BrPA and Azaserine treatments significantly impacted mESCs as they transitioned from epiblast-like to mesoderm stages in the directed differentiation system in vitro, while inhibition of late-stage glycolysis with YZ9 had no effect **(Figure S2D)**. For the treatment duration of our epiblast-to-mesoderm transition (day 1-4), mESC colonies were able to grow in size over time, and maintained significantly high levels of pluripotency genes including *Sox2, Pou5f1, Rex1* and *Klf2* while downregulating mesoderm genes including *T, Eomes, Mesp1*, and *Pdgfra* **(Figure 2F, G)**. Treatment duration targeting cells that had reached the mesodermal state (day 3-4) did not yield any phenotypic differences, and mesoderm-differentiated cells continued to blanket the plates on which they were cultured, similar to control conditions **(Figure 2F)**. However, these mesoderm-like cells showed significant *T, Eomes*, and *Mesp1* downregulation with reciprocally high *Sox2* and *Klf2* **(Figure 2G)**. These results further support our hypothesis that the inhibition of global glucose metabolism, as well as HBP, prevents cells from acquiring or maintaining a stable mesodermal identity. Additionally, we found that HBP-specific genes – *Gfat, Gnpnat1, Uap1*, and *Oga*– were also downregulated upon 2-DG + BrPA and Azaserine treatments (**Figures 2H, S2E)**.

FGF signalling has been known to play a vital role in mouse gastrulation for the specification of mesoderm at the posterior primitive streak, and has been linked with glycolytic activity in the presomitic mesoderm of developing chick embryos (Ciruna and Rossant, 2001; Oginuma et al., 2017). To test this relationship during mouse early mesoderm transition, we additionally treated mESCs with the FGF inhibitor PD0325901 (PD) during their directed-mesoderm differentiation, revealing overlapping phenotypes between FGF and glycolysis inhibited groups, and significant downregulation of mesodermal genes and upregulation of pluripotency genes **(Figures 2F, G)**. Interestingly, HBP-generated metabolites are known to play a role in FGF signal transduction through various post-translational modifications (Ghabrial, 2012; Kraushaar et al., 2012). The overlapping outcomes in 2-DG + BrPA, Azaserine, and PD treatments during embryo development and in vitro differentiation suggest an underlying relationship between glucose metabolism and FGF signalling in driving mesoderm specification and progression during mouse gastrulation.

### Glucose uptake is linked to an EMT programme

As high glycolytic activity was observed within the epiblast cells that preceded PS entry and mesoderm specification (Figure 1), we next investigated whether this regionalized phenotype was indicative of a metabolic role for the epithelial-to-mesenchymal transition (EMT) in mouse gastrulation. Strikingly, we found that the peak of Glut1 protein expression within the epiblast is specifically localized to cells adjacent to an intact basement membrane, as indicated by Laminin expression in sagittal, frontal, and transverse sections of mouse gastrulas **(Figure 3A, S3A)**. These high Glut1-expressing cells showed strong co-expression of Mmp14 protein, an extracellular matrix-degrading matrix metalloproteinase **(Figure 3B)**. The striking co-localization of this enzyme with Glut1 suggests that glucose uptake might be associated with breakdown of the basement membrane, a necessary step in EMT that would permit cell ingression into the PS. Indeed, embryos treated with 2-DG + BrPA or Azaserine were unable to break down the basement membrane in a proximal-to-distal manner, whereas embryos cultured with inhibitors of late-stage glycolysis, ATP synthase, lactate dehydrogenase, and the PPP and were able to perforate the ECM **(Figure 3C)**.

**Figure 3.**
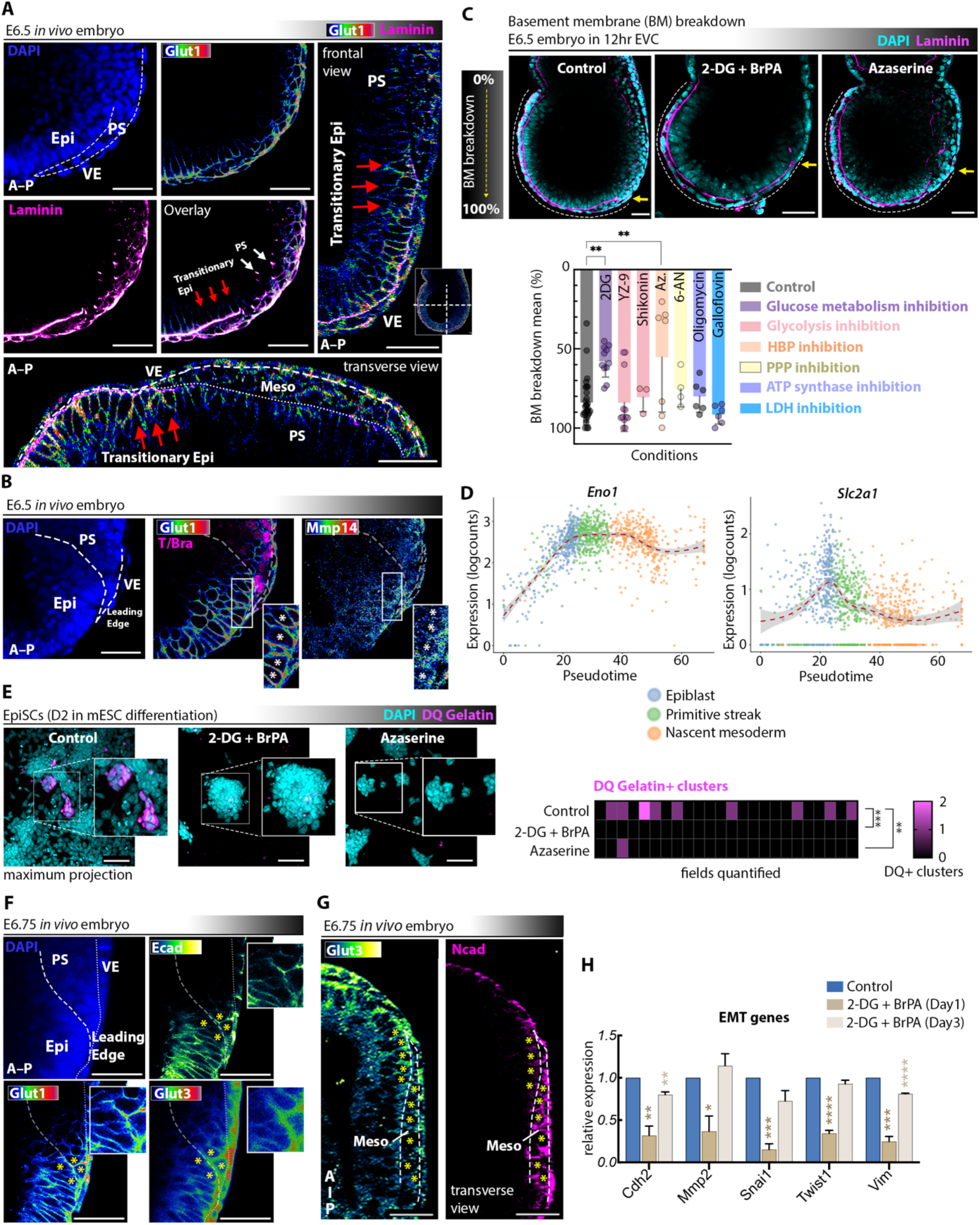
Glucose uptake is linked to an EMT programme. **(A)** Epiblast (Epi) Glut1 (heatmap intensity colours used via Fiji’s LUT) expression co-localises to regions of intact basement membrane identified via Laminin (magenta) staining (n=28 embryos), as well as regions of **(B)** high Mmp14 (heatmap intensity colours used via Fiji’s LUT) activity (n=3 embryos), particularly in Epi cells bordering the PS (white asterisks). Scale bars represent 40µm. **(C)** 2-DG + BrPA and Azaserine-treated embryos show a delay in their BM distal breakdown lengths, identified with Laminin (magenta) staining, while embryos cultured with other metabolic inhibitors show no such defect: Control (n=30), 2-DG + BrPA (n=11), YZ9 (n=12), Shikonin (n=3), Azaserine (n=7), 6-AN (n=4), Oligomycin (n=6), Galloflavin (n=6). Plots show mean ± SEM. Two-tailed parametric t-test. **P < 0.01. Scale bars represent 40µm. **(D)** Principal curves of glycolytic genes over pseudo-time (Epiblast to Primitive Streak to Nascent Mesoderm states) in gastrulating *in vivo* mouse embryos. **(E)** Representative outcome of an *in vitro* EMT assay with DQ Gelatin (magenta), applied on Epiblast-like Stem Cell (EpiSC) differentiation stage (Day 2, refer to Figure 2F). Quantifications show the number of DQ+ clusters identified in each imaging field (n=25 fields quantified over 2 independent experimental replicate). Two-tailed parametric t-test. **P < 0.01, ***P < 0.001 **(F)** Z-section of a posterior MS stage embryo showing Glut1/3 co-localizations to Cadherins, especially within epiblast and PS cell neighbours (ECad, white asterisk), and **(G)** within migratory mesoderm (NCad). Heatmap intensity colours used via Fiji’s LUT. Scale bars represent 40µm. **(H)** qPCR results of directed-differentiation experiments, show a downregulation of key EMT transcripts upon 2-DG + BrPA treatment across 3 experimental replicates. Plots show mean ± SEM. Two-tailed parametric t-test. *P < 0.05, **P < 0.01, ***P < 0.001, ****P < 0.0001.

To confirm that these metabolic changes occur during the epiblast to mesoderm transition *in vivo*, we analysed single-cell RNA sequencing (scRNAseq) datasets from E6.5, 7.0, 7.5, 8.0 and 8.5 stages from in vivo mouse embryos (Pijuan-Sala et al., 2019) and generated trajectory inference analyses of post-implantation epiblast, primitive streak and nascent mesoderm stages to model the transcriptional dynamics of the EMT process. This analysis identified shifts in gene expression of rate-limiting glycolytic enzymes, such as *Eno1, Pgam1, Gapdh, Ldha*, and *Ldhb*, as well as *Slc2a1*, and *Slc16a3* which were upregulated at the onset of EMT **(Figures 3D, S3B)**. In contrast, components of the Krebs Cycle, such as *Aco2, Mdh2, Sdha, Sdhb*, displayed a mostly static pattern of gene expression **(Figure S3B)**.

We next employed an *in vitro* EMT assay on directed-differentiated mesoderm using GFP-tagged DQ-Gelatin, a substrate that reveals bright fluorescence signal at sites of proteolytic digestion (Mook et al., 2003). In differentiating Epi-like cell colonies, the presence of many GFP+ cell clusters indicated regions on the plate where cells were actively undergoing a mesenchymal transition via the local release of MMPs (Mook et al., 2003) **(Figure 3E)**. As FGF is a known regulator of EMT in the posterior epiblast and is required for correct establishment of the PS in mouse development (Ciruna and Rossant, 2001), we treated cells with PD as a positive control for impaired proteolytic activity. As expected, PD-treated cells were devoid of GFP+ cell clusters **(Figure S3C)**. Likewise, treatment with 2-DG + BrPA or Azaserine also resulted in an absence of GFP signal, indicating that aberrant glucose metabolism functionally impairs EMT **(Figure 3E)**.

To further consolidate these findings, we analysed *in utero* dissected embryos for the presence of additional canonical EMT markers that might co-localise with glucose transporter (Glut1/Glut3) activity. Notably, we found both Glut1 and Glut3 protein expression to be specifically upregulated within the leading edge of PS and transitionary epiblast cells where E-Cadherin (ECad) expression is downregulated **(Figure 3F)**. Moreover, within the mesoderm wings where high N-Cadherin (NCad) expression persists, we also found high Glut3 expression **(Figure 3G)**. This points towards a role of glucose metabolism in cell adhesion during the EMT process, and suggests that enhanced glucose uptake spans the fate transition process of EMT **(Figure 3E, F)**. Further, qPCR analyses in mESC-differentiated mesoderm showed a significant reduction of distinct EMT genes upon 2-DG + BrPA treatment, including *Cdh2* (NCad), *Mmp2, Snai1, Twist1*, and *Vim* **(Figure 3H)**. Altogether, these findings reveal that the mouse gastrula’s EMT process is tightly linked to glucose metabolism, demonstrating how intracellular shifts in glycolytic activity can directly modulate cellular and molecular regulatory networks to control fate decisions.

### Glucose metabolism occurs upstream of FGF activity in the epiblast

FGF signalling plays an essential role for the specification of mesoderm and the coordination of gastrulation cell movements, with PS- and VE-expressed FGF8 acting non-autonomously on FGFR1+ embryonic epiblast (Ciruna and Rossant, 2001; Ciruna et al., 1997; Sun et al., 1999). Interestingly, various studies have shown that this signalling pathway is impaired when blocking protein-modifying enzymes that make use of substrates generated through glucose metabolism (Garcia-Garcia and Anderson, 2003; Shimokawa et al., 2011). Having demonstrated that the mouse embryo undergoes dramatic metabolic changes throughout the course of gastrulation, we wanted to understand what consequence the wave of glucose uptake might have for FGF signalling. Using immunofluorescence labelling, we first found high expression of diphosphorylated ERK (dpERK), a direct FGF signalling target, within transitioning epiblast cells and the newly formed mesodermal layer, but not within the PS itself **(Figure 4A)**. Notably, epiblast Glut1 expression was more anterior and more distal to dpERK signal, indicating that FGF induction might temporally follow glucose uptake within the epiblast **(Figure 4A)**. To understand the functional relevance of this phenotype, we treated embryos with 2-DG + BrPA and found that dpERK expression was absent upon glycolysis inhibition **(Figure 4A)**. Additionally, gastrulas cultured for 12hr *ex vivo* under 2-DG + BrPA and PD treatments were both unable to develop past the LS stage of gastrulation **(Figure 4B)**. These results suggest that our observed wave of glycolytic activity in the epiblast might be coupled to FGF signalling activity within the gastrulating embryo.

**Figure 4.**
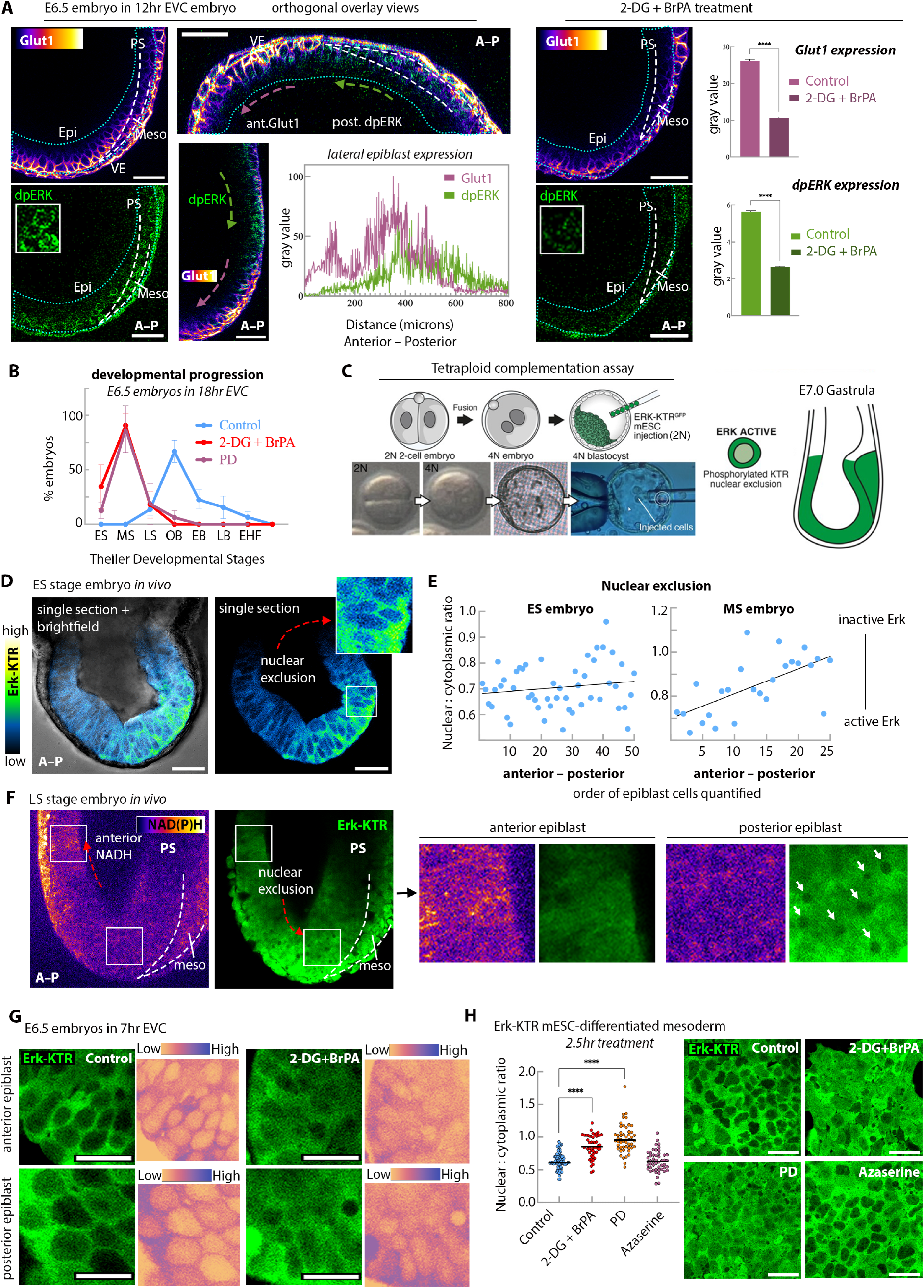
Glucose metabolism occurs upstream of FGF activity in the epiblast. **Left:** Single Z-sections of sagittal, frontal, and transverse views of control mouse embryo’s epiblast (green-dotted line) show that Glut1 expression (heatmap intensity colours used via Fiji’s LUT) is anterior and distal to dpERK expression (green). Plot profile shows expression intensities in the epiblast across the anterior-posterior axis of the orthogonal image above. **Right:** 2-DG + BrPA treatment abrogates Glut1 and dpERK activity in cells within the epiblast (green-dotted line): Control (n=3 embryos), 2-DG + BrPA (n=4 embryos). Scale bars represent 40µm. Two-tailed parametric t-test. ****P < 0.0001. Plots show mean ± SEM. Two-tailed parametric t-test. (B) Mouse gastrulas cultured with 2-DG + BrPA or PD for 18h *ex vivo* show equivalent developmental delay: Control (n=21), 2-DG + BrPA (n=24), PD (n=16). Plots show mean ± SEM. Two-tailed parametric t-test. **(C)** Schematic of tetraploid complementation assay, generating embryos where all cells of the embryo proper are derived from Erk-KTR^GFP^ mESCs. **(D)** Erk-KTR^GFP^ signal (heatmap intensity colours) from live-imaging allows for **(E)** quantifications of Erk activity by calculating nuclear-to-cytoplasmic (N:C) ratio in the epiblast tissue. As development proceeds, Erk activity becomes graded across the anterior-posterior axis. Plots show epiblast N:C ratio in representative ES (R^2^ = 0.24) and MS (R^2^ = 0.40) embryos generated by tetraploid complementation. Scale bars represent 40µm. **(F)** Multi-photon live imaging confirms that glycolysis (NAD(P)H activity) occurs anterior to Erk activation in the epiblast (nuclear-excluded regions). n=3 embryos. Zoom insets from anterior and posterior epiblast regions shown on the right. For NAD(P)H signal, heatmap intensity colours used via Fiji’s LUT. **G)** 2-DG + BrPA treatment of a representative MS gastrula stage embryo (generated by tetraploid complementation) decreases the strength of the nuclear-excluded phenotype of Erk activity in anterior and posterior epiblast regions. Heatmap intensity colours used on the right for better visualization. Scale bars represent 20µm. **(H)** Short 2-DG + BrPA treatment (2.5h) of mesoderm-differentiated Erk-KTR^GFP^ cells at day 4 shows a significant reduction in Erk activity (n≥50 cells per group quantified). Plots show mean ± SEM. Two-tailed parametric t-test. ****P < 0.0001. Scale bars represent 40µm.

In order to obtain a real-time, endogenous readout of FGF activity, we performed a tetraploid complementation assay (Nagy et al., 1993; Robertson et al., 1986) to generate “all mESC derived” epiblasts from ERK Kinase Translocation Reporter (KTR^GFP^) mESCs, which provides a quantitative read-out for FGF signalling activity through nuclear to cytoplasmic shuttling of the GFP tag (Pokrass et al., 2020; Simon et al., 2020) **(Figures 4C, S4A)**. Live imaging of these ERK-KTR^GFP^ tetraploid-embryos revealed a graded patterning of antero-posterior ERK activity within the epiblast **(Figures 4D, E, S4A)**, similar to our previously defined glucose uptake (see Figure 1). Further, live imaging of these embryos through multiphoton microscopy allowed us to view ERK-KTR^GFP^ alongside NAD(P)H activity. Notably, NAD(P)H signal is skewed anteriorly to the site of active ERK, supporting our hypothesis that the wave of glucose metabolism precedes FGF signalling dynamics within the gastrulating embryo **(Figure 4F)**. Importantly, when ERK-KTR^GFP^ embryos are subject to 2-DG + BrPA treatment, cells in the anterior and posterior epiblast domains both showed a decrease in ERK signal **(Figure 4G)**. Collectively, these findings led us to hypothesize that the wave of glucose uptake in epiblast tissue may be a prerequisite for functional FGF-mediated signal transduction.

To explore whether this relationship extends beyond the epiblast, we differentiated ERK-KTR^GFP^ mESCs to mesoderm *in vitro* and inhibited glucose metabolism (2-DG + BrPA), HBP (Azaserine), and FGF signalling (PD). We performed inhibitor treatments for only a brief 2.5h duration at D4 of differentiation, when cells have already reached a stable-mesoderm state, to avoid confounding effects of these perturbations on earlier differentiation steps that we previously showed to be dependent on these pathways (Figures 2F, G). Live-imaging of these cells revealed significant ERK inactivation upon short 2-DG + BrPA and PD treatments, but not in Azaserine groups **(Figure 4H)**. This excitingly suggested to us that glucose metabolism may be differentially regulated in epiblast vs. mesodermal tissue. In the same way that gastrulation is understood to be an integration of various processes, so too might we expect the relationship between glucose and FGF, whether mediated through HBP or late-stage glycolysis, to look different between various compartments of the developing gastrula.

### Late-stage glycolysis links with FGF signalling in the newly formed mesoderm

The FGF-inhibited phenotype of mouse gastrulas is widely recognizable for its posterior build-up of cells in the primitive streak, a result that is often attributed to a migratory failure of mesendoderm-progenitor exit from streak (Ciruna and Rossant, 2001; Sun et al., 1999) or from epiblast buckling due to an impaired mode of cell ingression (Najera and Weijer, 2023; Yamaguchi et al., 1994) **(Figure S4C)**. Despite our data supporting a close interaction between glucose metabolism and FGF signalling in epiblast tissue, 2-DG + BrPA and Azaserine-treated embryos do not recapitulate the impaired morphological PS phenotypes of PD-treated embryos **(Figures S4C)**. In a previous study of chick mid-gestation development, FGF signalling was shown to regulate the transcription of rate-limiting late-stage glycolytic enzymes in presomitic mesoderm (Oginuma et al., 2017). These glycolytic enzymes function by metabolizing the glucose-derived carbons that bypass HBP, and make up the segment of late-stage glycolysis that can be inhibited using YZ9 **(Figure 2A)**. This led us to believe that the critical function of HBP for gastrulation events might be limited to guiding cell fate transitions and EMT progression, and that HBP does not serve a critical function for cells once they have ingressed into the streak and become specified towards a mesoderm. Instead, these late-stage glycolytic enzymes may become important as soon as cells adopt mesodermal fate and begin their migration away from the midline of the embryo **(Figure 2E**).

To explore whether FGF signalling might regulate late-stage glycolysis in mesoderm-specified cells, we returned to our YZ9 treated embryos for a closer examination of their developmental phenotypes. Our earlier experiments showed no specific effect for mesoderm specification and PS elongation after YZ9 treatment **(Figures 2C, S2D)**. But after quantifying the PS surface area in sagittal sections of YZ9 and PD treated gastrulas, we found a significant increase in the PS size of these embryos **(Figures S4C, 5A)**. Subsequent immunolabeling of phospho-Histone H3 (to quantify cells undergoing mitosis) indeed revealed a marked increase in the number of dividing cells within the PS of PD as well as YZ9 treated embryos **(Figure 5B-D)**.

**Figure 5.**
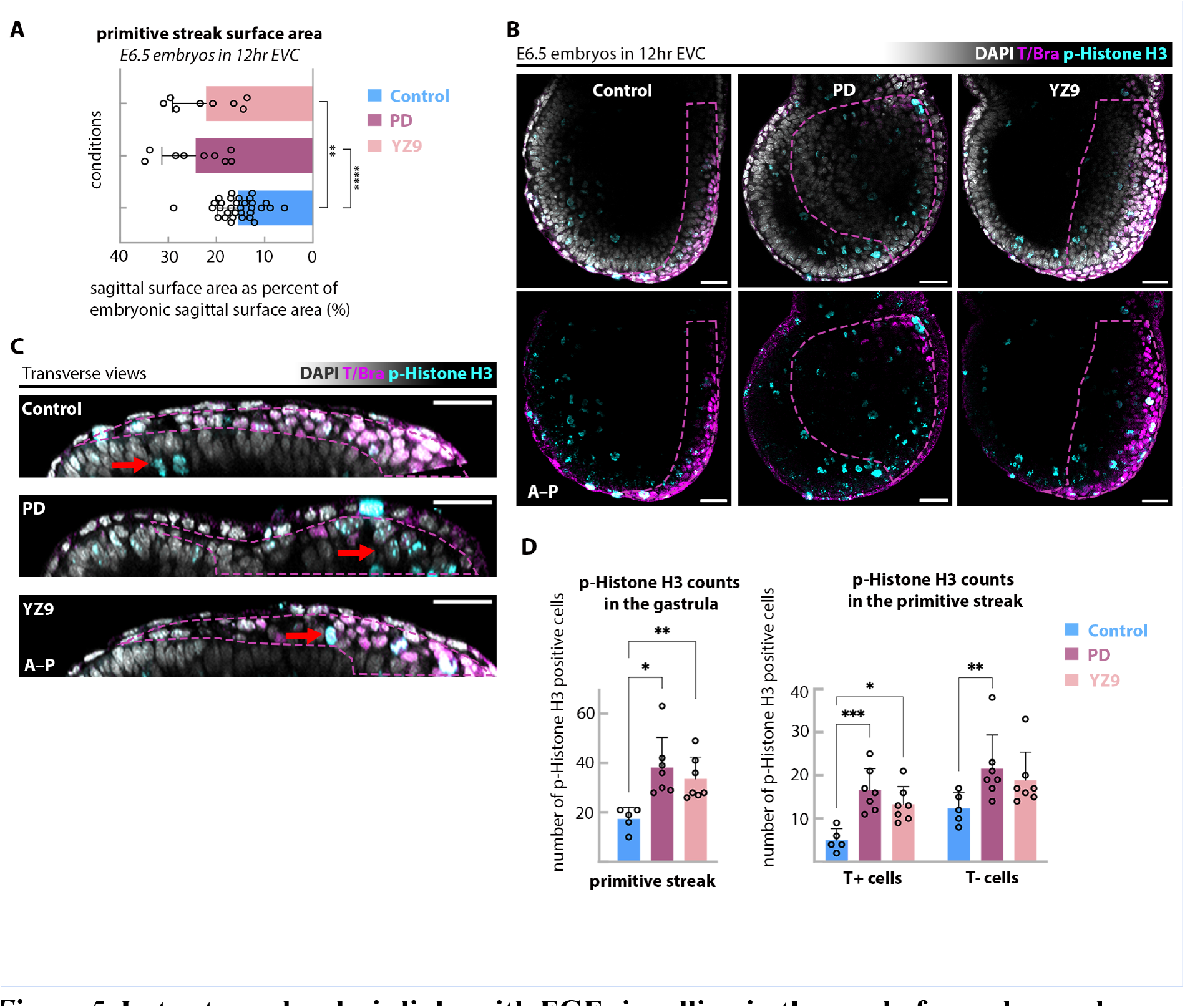
Late-stage glycolysis links with FGF signalling in the newly formed mesoderm. **(A)** PS surface areas calculated from mid-embryo sagittal Z-sections show a significant size increase in PD (n=9 embryos) and YZ9 (n=8 embryos) treated groups compared to control (n=32 embryos). Plots show mean ± SEM. Two-tailed parametric t-test. **P < 0.01, ****P < 0.0001. **(B)** Representative single Z-sections show that PS surface areas (marked by T/Bra, in magenta) are expanded in PD & YZ9 treated embryos (magenta-dotted regions), with high phospho-Histone H3 (cyan) expression in those regions: Control (n=7), PD (n=6), YZ9 (n=7). Scale bars represent 40µm. **(C)** Transverse sections confirm expanded PS phenotype and phospho-Histone H3 localizations in the PS (red arrows). Scale bars represent 40µm. **(D)** Phospho-Histone H3 counts in the PS (indicated via morphology or T expression) are significantly increased in PD and YZ9 treated embryos. Plots show mean ± SEM. Two-tailed parametric t-test. *P < 0.05, **P < 0.01,***P < 0.001.

As YZ9 treated embryos more closely recapitulate the FGF-inhibited phenotype, enhanced FGF signalling might play a role in regulating glycolysis within mesodermal cells, specifically at late-stage glycolytic steps of glucose metabolism. This seemingly mesoderm-specific mechanism would support observations in chick development (Oginuma et al., 2017).

### Glucose metabolism is necessary for proper mesoderm migration

The phenotypic similarity between YZ9 and PD treated embryos (Figures S4C, 5A-D), combined with high levels of glycolytic activity in mesenchymal mesoderm (as shown by Glut1/3 expression, NAD(P)H autofluorescence, and 2-NBDG localization; Figures 1C-J) suggest that late-stage glycolysis might have a functional role in mesoderm-specific cellular behaviour, such as motility. To assay cellular characteristics related to mesodermal migration, we first analysed lateral mesoderm lengths in mouse gastrulas, which revealed that 2-DG + BrPA, YZ9, and PD treatments yielded significantly shorter lateral mesoderm lengths relative to the embryos’ width **(Figure 6A)**. This finding was not observed in Azaserine-treated embryos, suggesting that HBP may not be required for the migration of mesodermal cells that have already exited the streak **(Figure 6A)**. To better understand how late-stage glycolysis might control migration dynamics, we isolated mesoderm explants from E7.25/5 LS/no bud stage mouse embryos (Burdsal et al., 1993; Levak-Svajger et al., 1969) from a H2B-GFP:Tcf-LEF reporter mice (Ferrer-Vaquer et al., 2010) to confirm mesoderm-specific isolation, and cultured these H2B-GFP:Tcf-LEF mesoderm explants on fibronectin-coated dishes for 24h **(Figure 6B)**. We first found a significant and specific decrease in the surface area of 2-DG + BrPA treated mesoderm explants compared to other inhibitor treatments, reflecting the cells’ inability to migrate away from the explant centre **(Figure S5A)**. However, PD and YZ9 treatment groups did not show a significant difference. **(Figure S5A)**. To better understand this outcome, we employed time-lapse live imaging and quantified cell movements **(Figures 6C, S5B)**. This showed a significant increase in proliferation rate, and a significant decrease in cell displacement rate following PD and YZ9 treatment compared to control groups **(Figure 6C, D)**. These measurements indicate that the morphological increase in explant surface area must be due to increased cell divisions, causing the explant to expand in size. It is thus, possible, that the expanded PS sizes of PD and YZ9 treated embryos (Figure S4C, 5A-D) is a result of increased proliferation, as well as aberrant cell migration.

**Figure 6.**
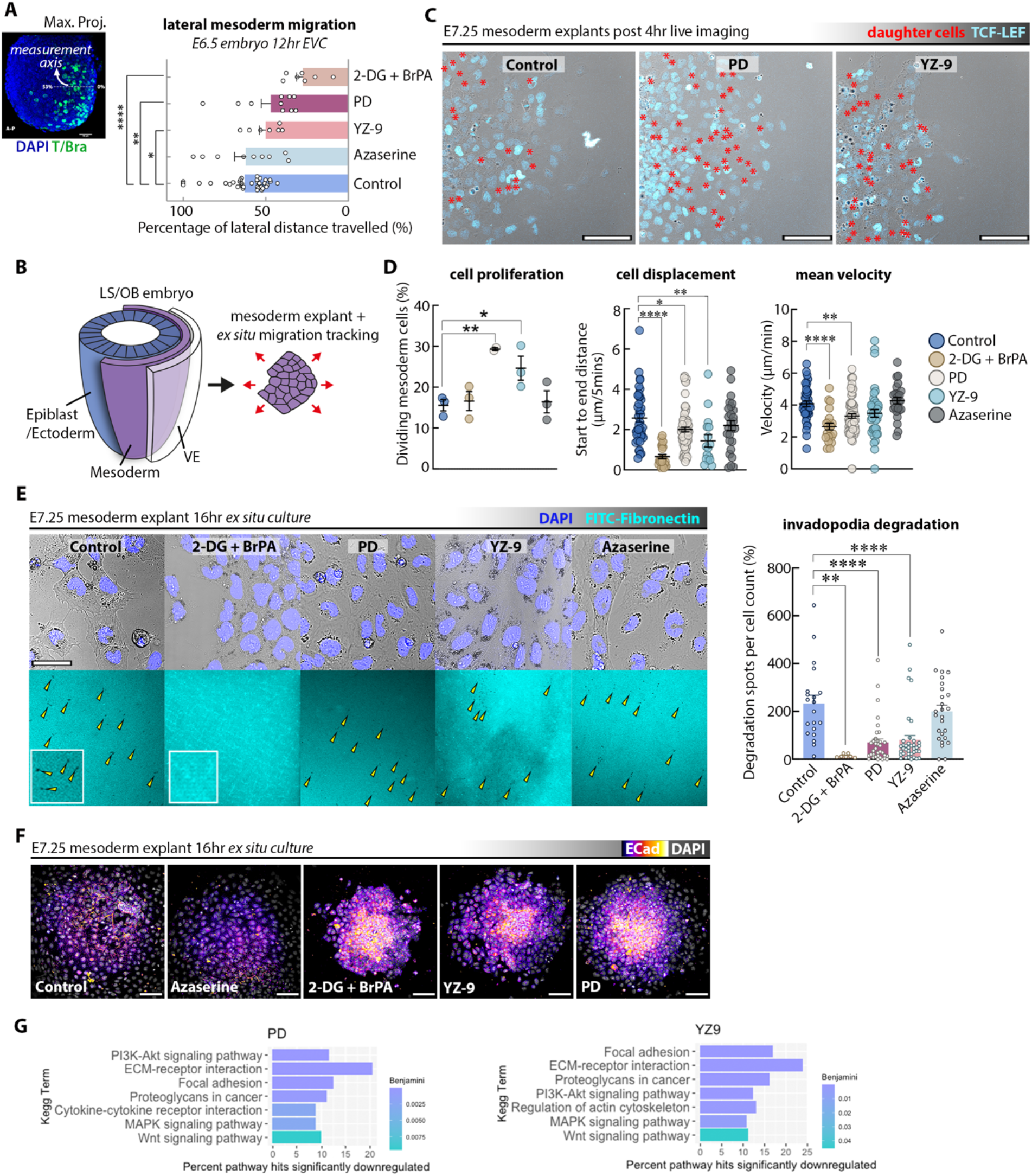
Glucose metabolism is necessary for proper mesoderm migration. **(A)** Average lengths of lateral mesodermal wings are significantly shorter in 2-DG + BrPA (n=7 embryos), YZ9 (n=7 embryos), and PD (n=10 embryos) treated embryos compared to control (n=38 embryos). Plots show mean ± SEM. Two-tailed parametric t-test. *P < 0.05, **P < 0.01,***P < 0.001. **(B)** Schematic of mesoderm explant isolations of embryos for subsequent live-imaging of *ex situ* migration dynamics. **(C)** Representative maximum projection images of Control, PD, and YZ9 treated mesodermal explants at the end of 4h live imaging, with red asterisks indicating daughter cells that have divided since the start of imaging. Reporter LEF-GFP signal (cyan) was used for nuclear segmentations and tracking. Scale bars represent 40µm. **(D)** Plots show migration readouts across 3 independent experimental replicates of explants generated from AIVIA cell-tracking software, showing that PD and YZ9 treated mesoderm have higher rates of proliferation and shorter migratory lengths compared to control. ‘Cell proliferation’ datapoints represent independent experiments. ‘Cell displacement’ and ‘mean velocity’ datapoints represent unique cell tracks. Plots show mean ± SEM. Two-tailed parametric t-test. *P < 0.05, **P < 0.01,***P < 0.001, ****P < 0.0001. Azaserine treatment shows no significant difference. **(E)** Invadapodia assay of mesoderm explants on FITC-Fibronectin-coated plates, showing impaired ECM substrate degradation in 2-DG + BrPA, YZ9, and PD treated explants (n=3 experimental replicates). Scale bars represent 100µm. Plots show mean ± SEM. Two-tailed parametric t-test. **P < 0.01, ***P < 0.001, ****P < 0.0001. Those treatment groups also resulted in **(F)** increased ECad expression (heatmap intensity colours used for better visualisation). Scale bars represent 100µm. **(G)** Pathway enrichment (KEGG) of the differentially expressed genes (DEGs) in PD and YZ9 treated mesoderm explants, specifically shows downregulated terms.

To functionally test whether or not these mesodermal cells have lost their full migratory capacity upon glycolytic inhibition, this time we explanted the mesoderm tissue isolated from wild-type embryos on FITC-Fibronectin, a fluorescent substrate that reveals fluorescent-devoid regions where ‘invasive’ mesenchymal cells have extended their protrusions to degrade the underlying extracellular matrix (ECM) (Artym et al., 2009) **(Figure 6E)**. Indeed, our invadopodia assay showed significantly lower amounts of substrate degradation in explants treated with 2-DG + BrPA, YZ9, and PD, indicating that both glycolysis and FGF-mediated ERK signalling affect proper mesoderm migration **(Figure 6E)**. Furthermore, immunolabelling of inhibitor-treated explants demonstrated a marked upregulation in adhesion marker E-Cad **(Figure 6E)**, while retaining mesenchymal marker N-Cad, as well as lateral mesodermal wing marker Eomes and Lef1 **(Figure S5C-D)** (Probst et al., 2021). As this is phenotype is consistent with previous literature on E-Cad overexpression in FGF mutant embryos (Ciruna and Rossant, 2001), the stark phenocopy with 2-DG + BrPA and YZ9 treated explants supports our hypothesis that FGF signalling may be tightly intertwined with glucose metabolism **(Figure 6F)**.

Finally, we performed RNA-sequencing on mesoderm explants treated with PD or YZ9 *ex situ*. Pathway enrichment analysis on mesoderm explants identified the differentially expressed genes revealing the downregulated KEGG (Kyoto Encyclopedia of genes and genomes) pathway terms which have direct roles with cell migration, such as “Focal Adhesion” (Choi et al., 2008; Gupton and Waterman-Storer, 2006; Parsons et al., 2010), as well as “ECM-Receptor interaction” and “MAPK Signalling Pathway” in both PD and YZ9 treated mesoderm explants **(Figure 6G)**. Additionally, differential GO (Gene Ontology) term analysis showed that most significantly downregulated processes were also similar between late-glycolysis and FGF inhibition, including essential aspects of mesoderm function such as focal adhesion, and integrin signalling **(Figure S5E)**. Collectively, these data support our hypothesis that an intimate connection between FGF signalling and late-glycolysis can regulate the processes that drive cell movements, allowing for the successful completion of mouse gastrulation during triploblastic development.

## Discussion

Mammalian embryos undergo an evolutionarily conserved switch towards glucose metabolism around the time of implantation, but the consequences of this metabolic shift have long been undefined. Here, we first characterised how glucose activity is coordinated across space and time in the mouse gastrula, discovering a stringent pattern of graded glucose uptake in pluripotent epiblast and migratory mesenchyme. Our further investigations revealed a tissue-specific and functionally distinct regulation of glucose metabolism, showing that HBP regulates cell fate transitions and EMT, while late-stage glycolysis drives cell migration. Importantly, we find that these events are tightly linked to FGF/ERK activity and suggest that a crosstalk between metabolism and morphogen signalling mechanistically regulates hallmark cellular and developmental programmes. Thus, these findings challenge a widely-held view of metabolism’s role in early embryogenesis to show that it is not only “life-sustaining”, but also “life-directing”.

HBP functions as a nutrient-sensing pathway and is at the centre of many cancer processes (Akella et al., 2019). Previous evidence demonstrates that its essential end-product, UDP-GlcNAc, is vital for the biosynthesis of heparan sulphate proteoglycan proteins (Hart et al., 2011; Hull et al., 2007; Mishra et al., 2011) which act as co-receptors for FGF signalling (Garcia-Garcia and Anderson, 2003). Additionally, UDP-GlcNAc is an essential substrate for various post-translational modifications, including GlcNAcylation and glycosylation events that are vital for cell fate transitions. As such, our observation that ERK signalling temporally follows glucose uptake in epiblast cells suggests a signalling transduction requirement of UDP-GlcNAC production. Indeed, a recent study using the mouse gastruloid model system also found that a glucose epimer, mannose, is required for mesoderm fate specification and axial elongation, and showed that FGF target genes such as Evt4 and Spry4 were downregulated following 2-DG treatment (Dingare et al., 2023). Together with this companion study, these findings provide further support for our data surrounding glucose-mediated cell fate transitions in mouse embryos, and underscore the importance of glucose metabolism for FGF signalling.

Mesendoderm patterned within the streak migrate anteriorly around the embryonic circumference (Saykali et al., 2019), and per our observations, show a high level of glucose uptake throughout this process. Intriguingly, our findings demonstrated that while HBP was dispensable for this event, glycolysis was required for embryos to complete this trilaminar transition. Moreover, we found that glycolysis-inhibited embryos exhibited a posterior expansion of cells within the PS that was reminiscent of the previously observed phenotype in FGF-mutant embryos (Ciruna and Rossant, 2001; Sun et al., 1999). Accordingly, our mesoderm explant migration assays revealed that this is due to cellular motility defects and aberrant proliferation. Our results show that glycolysis might act as the mechanistic link for FGF/ERK-mediated cell migration, suggesting that an impaired metabolism may serve as the underlying cause of FGF-abrogated morphogenesis.

We propose that these compartmentalised waves of glucose (HBP vs late-stage glycolysis) and FGF/ERK activity enables precise spatiotemporal regulation of key, stepwise morphogenetic events that drive gastrulation. Indeed, studies in both mouse and chick gastrulation suggest two temporal waves of ERK activity, first in cells prior to PS ingression, and then again during late-stage anterior migration (Lunn et al., 2007; McFann et al., 2022; Morgani et al., 2018). We hypothesize here that glucose metabolism acts as a conduit that allows ERK signalling to be co-opted for these distinct morphogenetic processes. The specifics of how FGF/ERK signalling interacts with distinct steps of glucose metabolism during mouse gastrulation, whether through post-transcriptional modifications by HBP (Czajewski and van Aalten, 2023; Tian et al., 2012), regulation of glycolytic transcripts (Oginuma et al., 2017), or a combination of both, remains to be explored.

The idea of metabolic guidance for developmental transitions and competence has gained popularity over the years, with each decade’s new set of experimental tools expanding our interpretation of the “gradient theory” first proposed in the first years of the 20^th^ century (Child, 1941). Though each are not without unique limitations, techniques like mass-spectrometry-based carbon tracing or the improved *ex vivo* embryo culture system have allowed us to press towards a more integrative view of embryogenesis (Chi et al., 2020). Collectively, the work we present here contributes to a growing body of work to support Child’s gradient theory of metabolite-guided morphogenesis, and describes, for the first time, how nutrient utilization occurs in concert with molecular signalling networks during mammalian gastrulation. At a time where one first acquires some semblance of organismal identity, “metabolic signalling” is a useful paradigm that allows us to come closer to an understanding of how the entire body plan is established.

Indeed, the best laid plans of mice [and humans] can often go awry, and our findings can provide important insights into the mechanisms behind developmental lethalities that arise from deregulated signals and perturbed metabolism. These results underscore the importance of metabolism for embryonic patterning and developmental success, and are highly applicable to other contexts where cellular transitions and migration take place, as in cancer biology and regeneration.

## Author contributions

D.C. and B.S. conceived the study. D.C. performed experiments and analysed the data on mouse embryos, tissue explants and stem cells. L.Z. performed mouse tetraploid complementation assays. AH and DC established and performed multiphoton live imaging of NAD(P)H in embryos and AH and VG helped with data interpretation. J.B performed analyses of RNA-seq datasets. A.L.C. provided critical discussions, provided illustrations presented and helped with manuscript writing. B.S. supervised the project, and wrote the manuscript together with D.C.

## Acknowledgements

We are grateful to Kaelyn Sumigray for the support in confocal microscopy, Zachary Smith and the Smith lab members for the support in microinjection and pseudopregrant mice preparation for tetraploid complementation assays. We thank all members of the Sozen lab for their helpful discussions throughout the project’s term, as well as Zachary Smith, Chaitanya Dingare and Ben Steventon for helpful feedback on the manuscript. AH is a NYSCF-Druckenmiller Fellow and is supported by The New York Stem Cell Foundation. VG is supported by an HHMI Scholar award (55108527), NIH grants number 1R01AR063663, 1R01AR067755 and the National Institute on Aging of the NIH under Award Number DP1AG066590. The Sozen lab is funded by NIH Early Innovators Award (DP2HD112040), the Richard and Susan Smith Family Foundation, the Yale Stem Cell Center Chen Innovation Award and Reprogrants. The content is solely the responsibility of the authors and does not necessarily represent the official views of the National Institutes of Health.

