## Supplementary Material for "A Spatiotemporal Compartmentalization of Glucose Metabolism Guides Mammalian Gastrulation Progression"

### Supplementary Figure 1

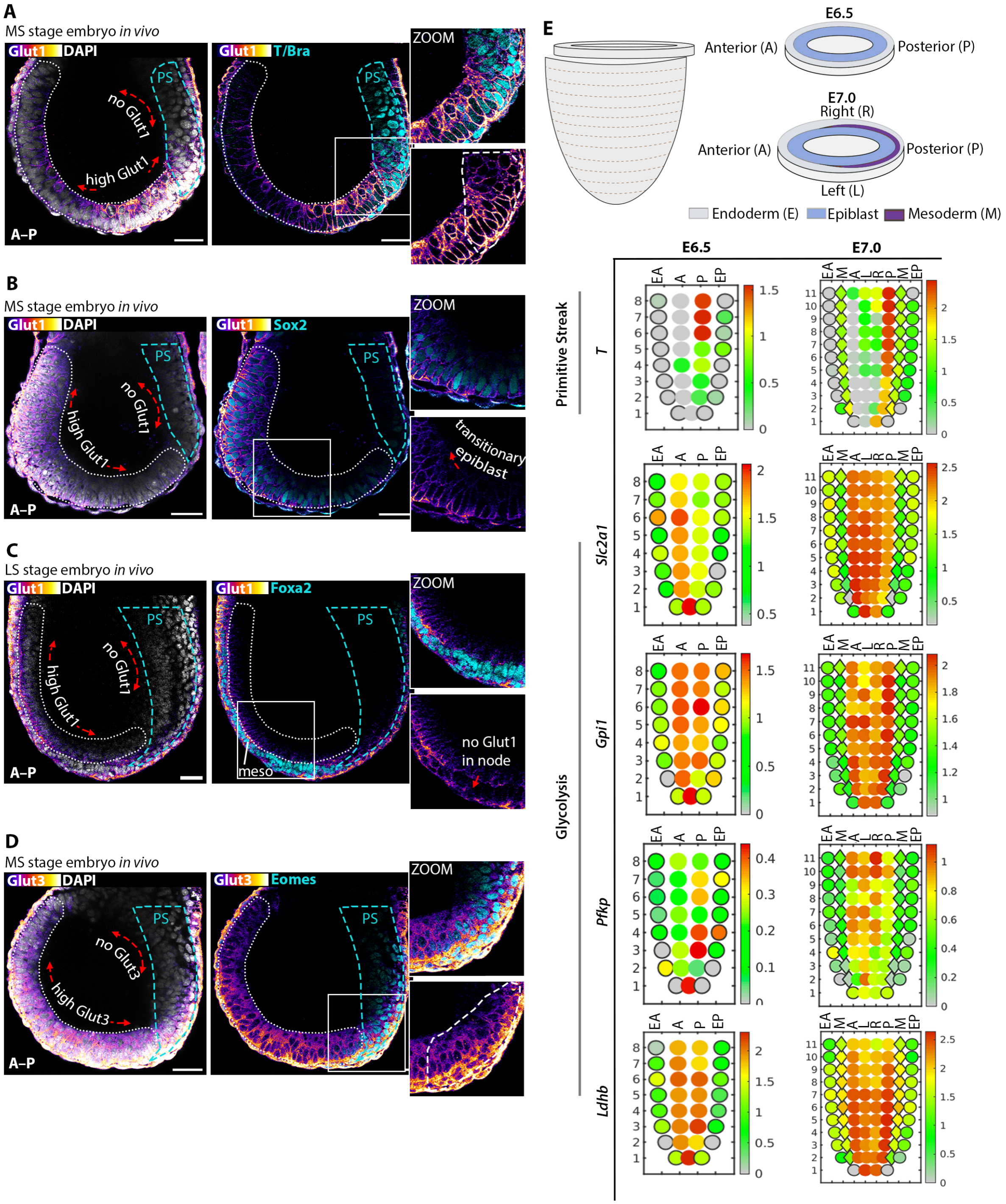

***Supplementary Figure 1.*** **(A-D)** Representative Z-sections of Glut1 (A-C) and Glut3 (D) expression (heatmap intensity colours used via Fiji's LUT) in relation to cell-types of interest in mid (MS) or late streak (LS) stage mouse gastrulas. Zoom insets of the epiblast and PS border shown on right. A, Anterior; P, Posterior. ($n \geq 9$ embryos per staining condition) Scale bars represent 40µm. **(E) Top:** Schematic of mouse embryo transverse planes showing laser capture microdissection transcriptome spatial coordinates of **Bottom:** corn plots of primitive streak gene *T*, and glycolytic genes *Slc2a1, Gpi1, Pfkp, Ldhb*, generated by querying genes of interest from the online e-gastrulation Geo-seq database (Peng et al 2016). A, Anterior; P, Posterior; AE, Endoderm Anterior; EP, Endoderm Posterior; M, Mesoderm; L, Left; R, Right.

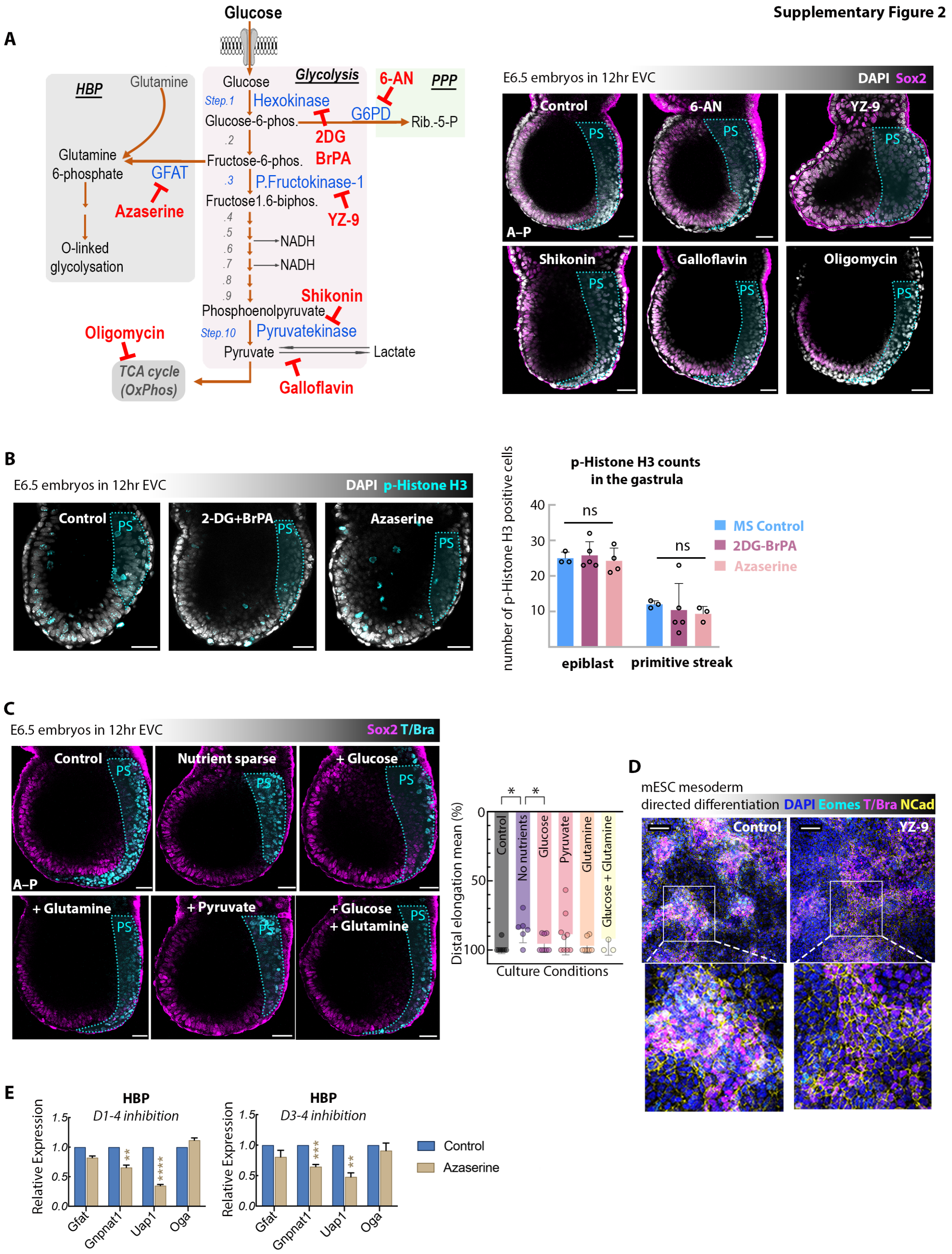

A
Glucose
HBP
Glutamine
Glutamine 6-phosphate
GFAT
Azaserine
O-linked glycolysation
Glycolysis
Step.1
Hexokinase
Glucose-6-phos.
2DG
BrPA
Fructose-6-phos.
P.Fructokinase-1
Fructose1.6-biphos.
YZ-9
NADH
NADH
Shikonin
Phosphoenolpyruvate
Step.10
Pyruvatekinase
Pyruvate
Lactate
Galloflavin
Oligomycin
TCA cycle (OxPhos)
6-AN
PPP
G6PD
Rib.-5-P
E6.5 embryos in 12hr EVC
DAPI Sox2
Control
6-AN
YZ-9
Shikonin
Galloflavin
Oligomycin
PS
A–P
B
E6.5 embryos in 12hr EVC
DAPI p-Histone H3
Control
2-DG+BrPA
Azaserine
p-Histone H3 counts in the gastrula
number of p-Histone H3 positive cells
ns
MS Control
2DG-BrPA
Azaserine
epiblast
primitive streak
C
E6.5 embryos in 12hr EVC
Sox2 T/Bra
Control
Nutrient sparse
+ Glucose
+ Glutamine
+ Pyruvate
+ Glucose + Glutamine
Distal elongation mean (%)
Control
No nutrients
Glucose
Pyruvate
Glutamine
Glucose + Glutamine
Culture Conditions
D
mESC mesoderm directed differentiation
DAPI Eomes T/Bra NCad
Control
YZ-9
E
HBP
D1-4 inhibition
D3-4 inhibition
Relative Expression
Gfat
Gnpnat1
Uap1
Oga
Control
Azaserine

***Supplementary Figure 2.*** **(A) Left:** Schematic of glucose metabolism and its three branches: grey = Hexosamine Biosynthetic Pathway (HBP); pink = core/late-stage glycolysis; green = Pentose Phosphate Pathway (PPP). Red text indicates chemical inhibitors and their metabolic targets in blue. **Right:** Representative images show mouse gastrula developmental outcomes following 12hr metabolic inhibitor treatment. A, Anterior; P, Posterior. Scale bars represent 40µm. **(B)** Representative Z-sections showing similar phospho-Histone H3 localizations (cyan) in 2-DG-BrPA and Azaserine treated embryos compared to control embryos at the MS stage of gastrulation. Scale bars represent 40µm. Plots show mean ± SEM. Two-tailed parametric t-test. **(C)** Mouse gastrulas cultured in nutrition-sparse media (free of glucose, pyruvate, and glutamine), and rescue conditions (selective reintroduction of the excluded nutrients) to functionally validate the specific effects of chemical inhibitors. A, Anterior; P, Posterior. Scale bars represent 40µm. Plots show mean ± SEM. *P < 0.05 Two-tailed parametric t-test. **(D)** Eomes (cyan), T (magenta), and NCad (yellow) expression remains unchanged for mesoderm-differentiated cells under YZ9 treatment at Day 4. Scale bar represents 100µm. 2 independent differentiation experiment. **(E)** qPCR analyses of directed differentiation experiments under Azaserine treatment, querying transcript changes in HBP genes. Plots show mean ± SEM. 3 experimental replicates. Two-tailed parametric t-test. **P < 0.01, ***P < 0.001, ****P < 0.0001.

### Supplementary Figure 3

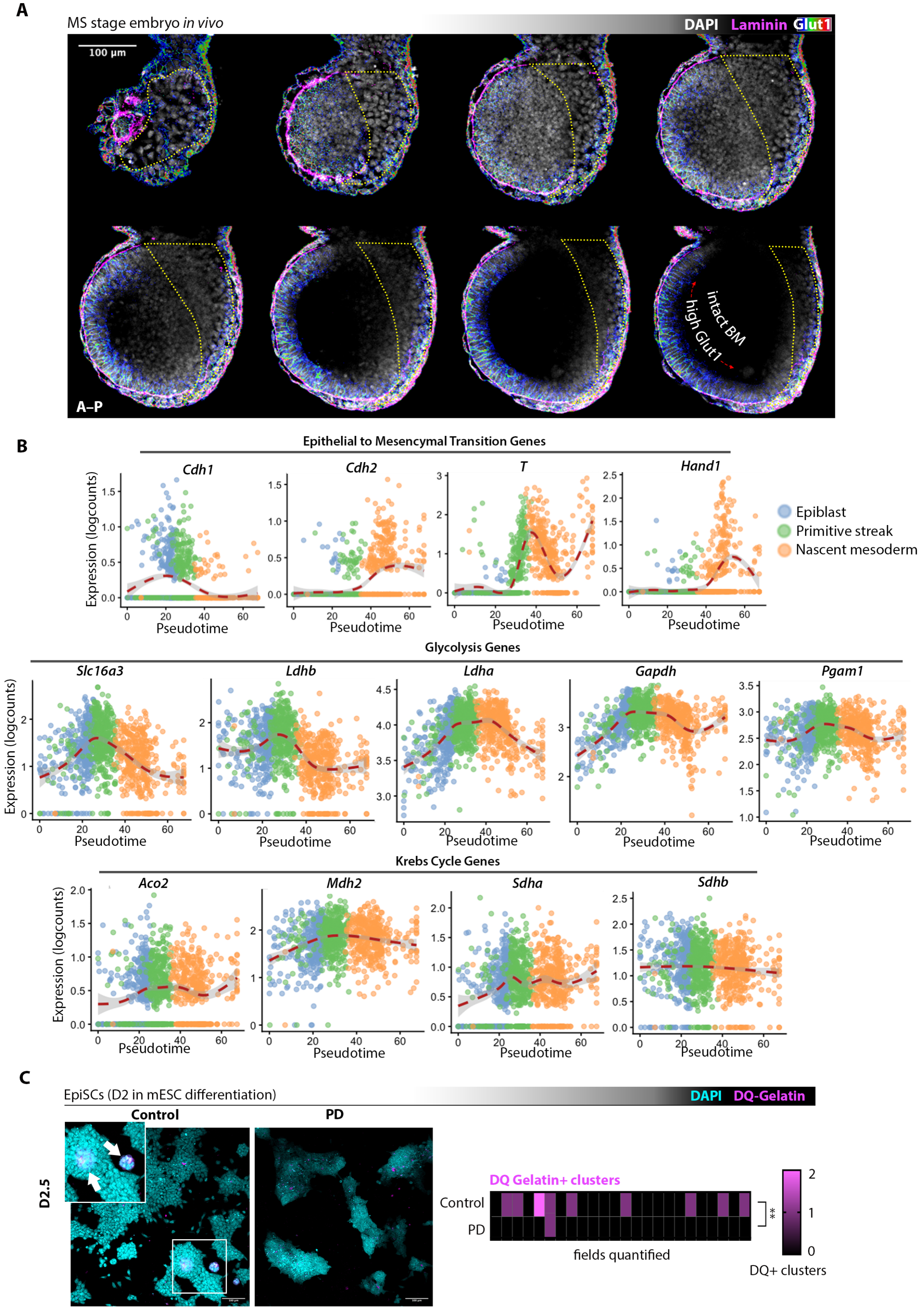

***Supplementary Figure 3.*** **(A)** Representative montage of sagittal Z-sections through the MS stage embryo shows epiblast Glut1 (heatmap intensity colours used via Fiji's LUT) expression co-localizing to regions of intact basement membrane, as identified via Laminin (magenta) stainings (n=28 embryos). A, Anterior; P, Posterior. Scale bar represents 100µm. **(B)** Principal curves of EMT, glycolysis, and Krebs Cycle genes over pseudo-time (Epiblast to Primitive Streak to Nascent Mesoderm states) in gastrulating *in vivo* mouse embryos. **(C) Left:** Representative outcome of a PD-treated *in vitro* EMT assay with DQ Gelatin (magenta). **Right:** Quantifications show the number of DQ+ clusters identified in each imaging field (n=25 fields quantified over 2 independent experimental replicates). Two-tailed parametric t-test. **P < 0.01.

**Supplementary Figure 4**

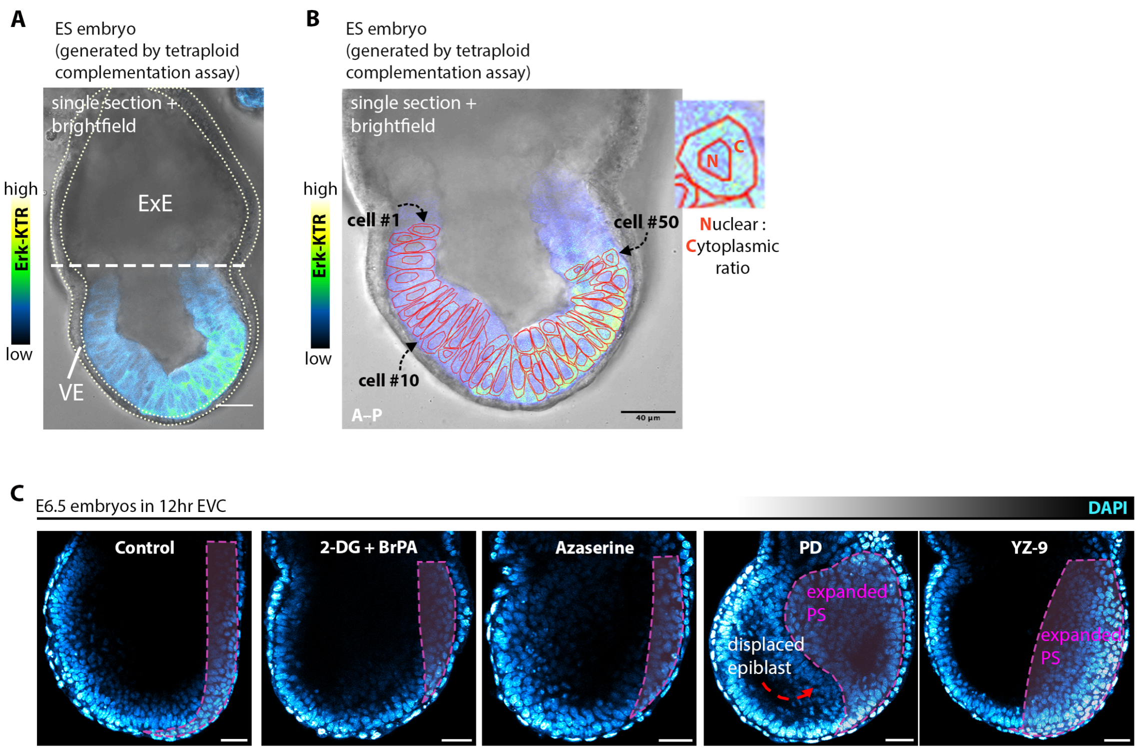


***Supplementary Figure 4.*** **(A)** Tetraploid complementation assay generates embryos where cells of the embryo proper are only derived from Erk-KTR$^{GFP}$ mESCs. Scale bar represents 40µm. **(B)** Nuclear and cytoplasmic manual segmentations of an ES stage embryo (generated by tetraploid complementation) were used to quantify nuclear-to-cytoplasmic ratios (N:C) of Erk-activity, quantified in order of anterior (A) to posterior (P) location in the epiblast. Scale bar represents 40µm. **(C)** Representative sagittal sections of mouse gastrulas treated with inhibitors (2-DG-BrPA, Azaserine, YZ9, or PD) for 12h *ex vivo*. PD treatment results in a build-up of cells within the primitive streak region, displacing epiblast cells in the process. Similar trend of posterior-increase observed under YZ9 treatment. A, Anterior; P, Posterior. Scale bar represents 40µm.

### Supplementary Figure 5

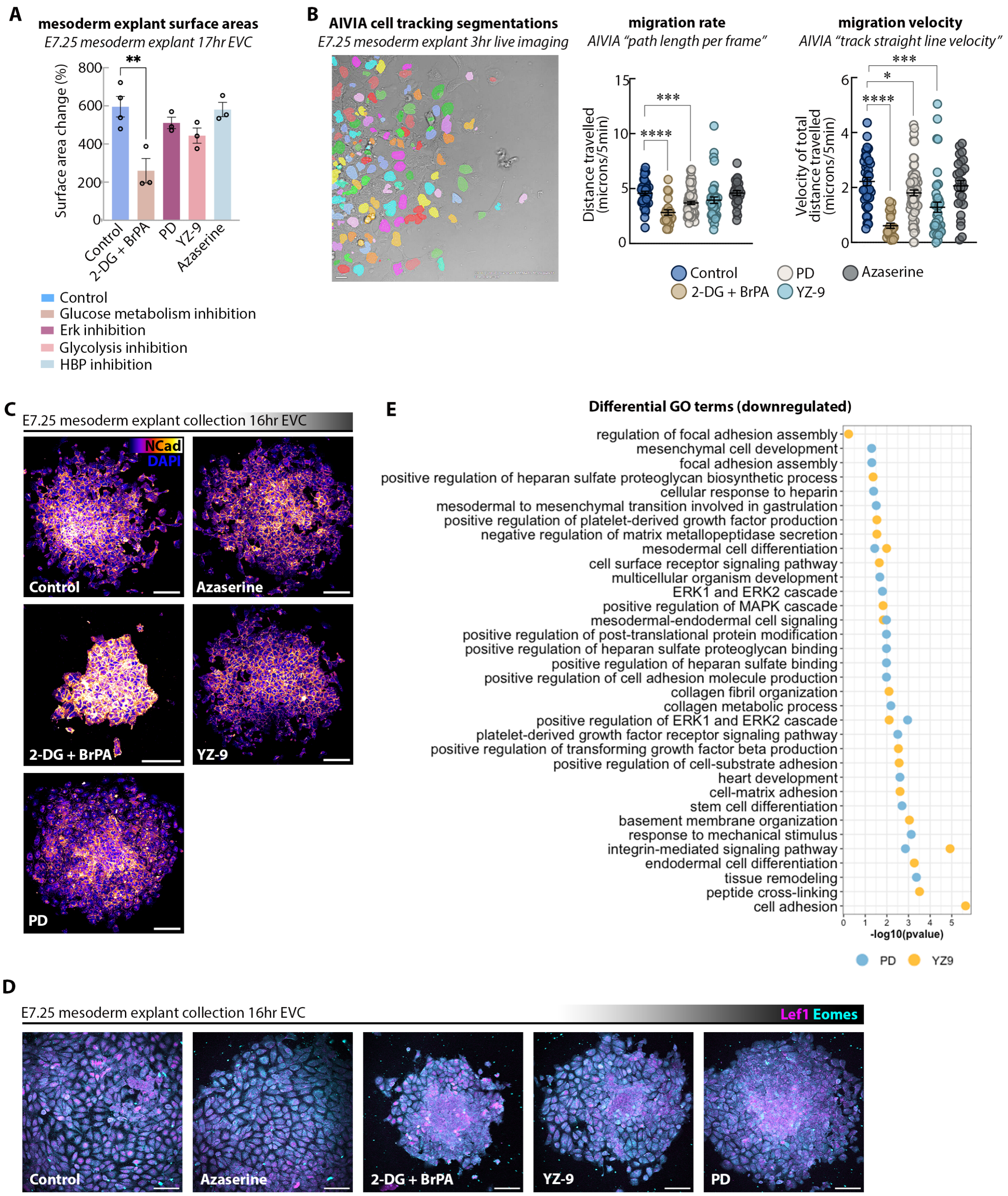

***Supplementary Figure 5.*** **(A)** Surface area changes of mesoderm explants following 17hr *ex situ culture* show that only 2-DG-BrPA treatment results in a significant size decrease compared to other groups. Plots show mean ± SEM. 3 biological replicates. Two-tailed parametric t-test. *P < 0.05, **P < 0.01, ***P < 0.001, ****P < 0.0001. **(B)** AIVIA "cell tracking" software allows for automated segmentations of live-imaging videos to track the migration rate (graph on the left) and velocity (graph on the right) of TCF/LEF-GFP mesodermal explants across different treatment groups. Individual datapoints represent unique cell tracks. Plots show mean ± SEM. Two-tailed parametric t-test. *P < 0.05, ***P < 0.001, ****P < 0.0001. **(C)** NCad expression (heatmap intensity colours used via Fiji's LUT) in mesoderm explants is retained across different treatment groups. Scale bar represents 40µm. **(D)** Lef1 (magenta) and Eomes (cyan) expression in mesoderm explants are consistently expressed across treatment groups (Azaserine, 2-DG-BrPA, YZ9, PD) after 16h. Scale bar represents 40µm. **(E)** GO biological process enrichment of downregulated genes in mesoderm explants after PD or YZ9 treatments.

### Methods

**Mouse embryo recovery.** Mice were maintained in accordance with national and international guidelines. All experimentation involving animal subjects was approved by the Institutional Animal Care and Use Committee at Yale School of Medicine and conducted following the approved animal handling protocol. All experimental mice were maintained in specific pathogen-free conditions on a 12–12 -hr light-dark cycle temperature-controlled facility with free access to water and food, and used from 6 weeks of age. All mice were bred to a mixed CD1 albino background. Valentina Greco (Yale University) provided the H2B-GFP:Tcf-LEF reporter mouse line. The experiments were not randomized. The investigators were not blinded to allocation during experiments and outcome assessment.

To collect *in vivo* embryos, 5-to-7 week-old female CD-1 mice were naturally mated with 12-to-24 week-old male CD-1 mice and sacrificed 6.5, 6.75, 7.0 or 7.25 days post coitum. Uteri were recovered and embryos were dissected from deciduae in DMEM medium containing 5% FBS, 20mM HEPES, and 1X Penicillin-Streptomycin (Gibco) warmed up to 37º. The sex of the embryos was not determined.

***Ex vivo* embryo culture.** Embryos were cultured in a 1:1 ratio of DMEM F/12 (Gibco) to rat serum (B4520 Envigo), with 1X Glutamax (Gibco), 0.2X Penicillin-Streptomycin (Gibco), and 0.2X MEM NEAA (Gibco) in 37°C at 20% O2 and 5% CO2. Media was equilibrated in 5% CO2 for ≥15mins prior to culture.

**Metabolism and signal modulation experiments.** Chemical inhibitor concentrations were administered as follows: 2mM 2-DG (Santa Cruz Biotechnology), 20µM BrPA (Santa Cruz Biotechnology), 10µM PD0325901 (StemCell Technologies), 200µM Galloflavin (Cayman Chemical), 100µM Oligomycin A (Santa Cruz Biotechnology), 5µM 6-Aminonicotinamide (Cayman Chemical), 10µM YZ9 (Cayman Chemical), 5µM Shikonin (Cayman Chemical), and 5µM Azaserine (Cayman Chemical). Embryos were treated for 7h, 12h, or 18h. Mesoderm explants were treated for 3h, 12h, or 16h. Stem cells were treated for the duration of the experiment as indicated in the figure captions.

**Nutrient-sparse and rescue experiments.** Nutrient sparse media was prepared using an Advanced DMEM/F-12 media devoid of D-glucose, L-serine, L-glutamine and sodium pyruvate (Caisson Laboratories) with the addition of 25mM sodium bicarbonate (Sigma), 0.2X Penicillin-Streptomycin (Gibco), and 0.2X MEM NEAA (Gibco). This was supplemented with the following nutrients, according to each experimental condition: 17mM glucose (Gibco), 0.5mM sodium pyruvate (Gibco), and/or 1X Glutamax (Gibco). All embryos were cultured in a 1:1 ratio of customised DMEM to rat serum (Envigo).

**Tetraploid complementation.** Mouse embryos at the two cell stage were recovered in KSOM medium by flushing the oviduct, from 5-6 weeks old CD1 females that were superovulated by injection of 10 IU of pregnant mares' serum gonadotropin (ProSpec) followed by 10 IU of human chorionic gonadotropin (Sigma) after 48 hours and were mated with CD1 males. Two cell stage embryos were then fused to induce tetraploidy using BTX Embryo Manipulation Electro Cell Fusion System. Fused embryos were transferred to advanced KSOM (Sigma) covered with mineral oil (FUJIFILM) and cultured to blastocyst stage until ESC injection. Each host tetraploid blastocyst was injected 10-15 mouse embryonic stem cells (mESCs), the injected blastocysts were transferred into uterus of the prepared D2.5 CD1 pseudopregnant surrogates (6 weeks old) which were plugged by vasectomized CD-1 male mice. These mESC-derived embryos were collected at the certain developmental timeline for subsequent experiments through uterine dissection (described above).

**Cell culture and in vitro differentiation assay.** All cells were cultured at 37°C in 20% O2 and 5% CO2 and passaged once they had reached 80% confluency. Cells were routinely tested for mycoplasma contamination by PCR.

mESCs were cultured on gelatinized tissue-culture–grade plates in FBS-containing DMEM medium with 2i/LIF (1μM MEK inhibitor PD0325901, 3μM GSK-3 inhibitor CHIR99021 and 10ng ml−1 LIF). FBS-containing DMEM (Gibco) medium comprised of: 18% inactivated FBS (Gibco), 1.2X Penicillin-Streptomycin (Gibco), 1.2X Glutamax (Gibco), 1.2X MEM NEAA (Gibco), 1.2mM Sodium Pyruvate (Gibco), and 120μM 2-Mercaptoethanol (Gibco).

For *in vitro* directed differentiation experiments, we followed the protocol as described previously in Gouti et. al. (2014) with some modifications. Cells were passaged (day 0) after reaching 80% confluency into 6-well culture plates or 8-well μ slides (Ibidi) at a density of 20k/1cm$^2$ in FBS-containing DMEM medium with 10ng/mL Fgf2 (R&D Systems). Cells were treated at day 1 with 10ng/mL Fgf2, at day 2 with 10ng/mL Fgf2 and 5μM CHIR99021 (StemCell Technologies), and at day 3 with 5μM CHIR99021.

**In situ mesoderm explant assay.** For mesoderm isolation from E7.25 mouse embryos, we followed a modified protocol as described in Burdsal et al. (1993). Embryos were collected as described above. Extraembryonic tissue proximal to the amnion was removed, and the cup-shaped embryo was transferred to a dissociation media consisting of 0.5% Trypsin EDTA (Gibco) and 2.5% Pancreatin (Thermo Scientific Chemicals) for 15min in 4º. Embryos were washed with collection media (described earlier) and mesoderm tissue was dissected using insect pins (Roboz) attached to syringes. Explants were transferred to fibronectin-coated 18-well μ slides (Ibidi) plates and left to adhere for 3-4hr in media containing a 1:4 ratio of rat serum to embryo culture media (described earlier), prior to downstream experiments and/or chemical inhibitor treatments. For live imaging of migration dynamics, explants were imaged every 5min up to 3h. For immunofluorescence staining, explants were cultured up to 17h and fixed in PBS containing 4% PFA for 20min. For RNA sequencing, explants were cultured to 27h then scraped off with sterile insect pins and flash-frozen.

**Glucose uptake assay.** Mouse embryos were collected at ES, MS, and LS gastrulation stages and cultured with 1mM 2-NBDG (Cayman Chemicals) for 2h. For multi-photon microscopy, embryos were immediately live-imaged. For confocal microscopy, embryos were fixed in PBS containing 4% PFA for 15min then immediately imaged after a 5min wash in PBS-T [PBS with 0.05% Tween-20].

**DQ gelatin assay.** Cells underwent mesoderm-directed differentiation on 8-well μ slides (80826 Ibidi) as previously described. On day 2.5 when cells are at the EpiSC stage, cells were cultured with differentiation medium including 50μg/mL DQ$^{TM}$ gelatin (Invitrogen) then fixed on day 3.5 in PBS containing 4% PFA for 20min, protected from light. Cells were incubated in blocking buffer (described below) with DAPI (overnight at 4ºC or 20min RT) prior to imaging.

**Invadopodia assay.** 18-well μ slides (Ibidi) were prepared with FITC-Fibronectin (Sigma-Aldrich) and 0.1% gelatin following the manufacturer's 2-day protocol and protected from light. Explants were cultured on these plates for 16h prior to fixation in PBS containing 4% PFA for

20min. Explants were incubated in blocking buffer (described below) with DAPI (overnight at 4ºC or 20min RT) prior to imaging.

**Immunofluorescence staining.** Samples were fixed in PBS containing 4% PFA for 20-45 min RT, or in methanol for 20min at 4ºC where stated. After fixation, samples were washed twice with PBS-T [PBS with 0.05% Tween-20] and permeabilised in PBS with 1mM Glycine and 0.3% Triton X-100 for 20-60 min at RT. Primary antibody incubations took place overnight at 4°C in blocking buffer [PBS containing 10% fetal bovine serum (FBS), 10% Tween-20]. Samples were washed twice with PBS-T prior to secondary antibody incubations at 4°C in blocking buffer. On the final day, samples were washed twice with PBS-T, then transferred into PBS-A droplets [PBS with 0.75% Bovine Albumin Fraction V (Gibco)] and covered with 30mm mineral oil (Sigma Aldrich) in 35mm glass-bottom dishes (MatTek) before confocal imaging. All PBS-T washes were done for 10min RT, and all RT incubations or washes took place on a rocking platform. All antibodies used in this study are listed in Supplementary Table 1.

**Supplementary Table 1.** Antibodies used in this paper

| Antigen | Host | Dilution | Cat. Number |
|---|---|---|---|
| Glut1 | Ms | 1:250 | MA1-37783 (Invitrogen) |
| Snai1 | Gt | 1:200 | AF3639 (R&D Systems) |
| Glut3 | Rb | 1:250 | MA532696 (Invitrogen) |
| NCad | Ms | 1:200 | MA1-91128 (Invitrogen) |
| Sox2 | Rt | 1:800 | 14-9811-82 (Invitrogen) |
| T/Bra | Gt | 1:500 or 1:200 | AF2085 (R&D Systems) |
| T/Bra | Rb | 1:250 | 81694S (Cell Signaling) |
| Lef1 | Rb | 1:400 | MA5-14966 (Invitrogen) |
| Laminin | Rb | 1:500 | PA1-16730 (Invitrogen) |
| Mmp14 | Rb | 1:250 | PA5-13183 (Invitrogen) |
| Ecad | Rt | 1:200 | 13-1900 (Invitrogen) |
| dpErk | Rb | 1:100 | 4377T (Cell Signaling) |
| P-Histone H3 | Rb | 1:500 | 441190G (Invitrogen) |
| Foxa2 | Rb | 1:200 | 8186S (BD Biosciences) |
| Eomes | Rt | 1:200 | 53-4875-80 (Invitrogen) |
| DAPI 405 | - | 1:500 | D3571 (Invitrogen) |
| anti-mouse Alexa Fluor 488 | Donkey | 1:500 | A21202 (Invitrogen) |
| anti-rabbit Alexa Fluor 488 | Donkey | 1:500 | A21206 (Invitrogen) |
| anti-rat Alexa Fluor 488 | Donkey | 1:500 | A21208 (Invitrogen) |
| anti-mouse Alexa Fluor 568 | Donkey | 1:500 | A10039 (Invitrogen) |
| anti-goat Alexa Fluor 568 | Donkey | 1:500 | A11057 (Invitrogen) |
| anti-rabbit Alexa Fluor 568 | Donkey | 1:500 | A10042 (Invitrogen) |
| anti-mouse Alexa Fluor 647 | Donkey | 1:500 | A31571 (Invitrogen) |
| anti-rabbit Alexa Fluor 647 | Donkey | 1:500 | A31573 (Invitrogen) |
| anti-goat Alexa Fluor 647 | Donkey | 1:500 | A21447 (Invitrogen) |
| anti-rat Alexa Fluor 647 | Donkey | 1:500 | A78947 (Invitrogen) |

**Image data acquisition and processing.** Samples were imaged with the Leica STELLARIS 5 microscope using a HC PL APO CS2 40x/1.10 water objective, a Z-spacing of 0.75 µm to 5µm and appropriate laser/filters for Alexa 405, Alexa 488, Alexa 546, and Alexa 633 or combinations thereof. To correct for fluorescence decay along the Z-axis during embryo imaging, 'Z-compensation by AOTF and PMT' was defined in a control embryo and applied across all experimental conditions during that imaging session, so that changes in laser power and gain across the Z-stack were equivalent across conditions, and only adjusted to each embryo's size. Raw data were processed using open-source image analysis software Fiji/ImageJ2 2.9.0 or AIVIA 10.5.1 AI Image Analysis Software and assembled in Photoshop 2021 22.3.1 (Adobe). Transverse views were generated from 1µm Z-spaced images using ImageJ's orthogonal viewer. Digital quantifications and immunofluorescence signal intensity graphs were obtained using Image J's plot profile measurements and visualised in GraphPad Prism9.5.0 software.

**Time-lapse live imaging.** Confocal time-lapse imaging of embryos, mesoderm explants, and mesoderm-differentiated cell cultures were performed using Leica STELLARIS 5 microscope using a 25x (HC FLUOTAR L 25x/0.95 W VISIR 0.17) or 40x (HC PL APO CS2 40x/1.10) water objective and appropriate laser/filters for Alexa 488, Alexa 546, and Alexa 633 or combinations thereof. Samples were imaged under a humidified chamber with 37 °C and 5% CO2. Explants were imaged at 5min intervals in 2µm Z-spaced planes for up to 4h on pre-treated ibidi dishes (Ibidi, USA). Images were processed using AIVIA, described below under 'image analysis'.

**Multiphoton live imaging for NAD(P)H autofluorescence.** Multicolour two-photon microscopy was used for live-embryo imaging of NAD(P)H dynamics. Embryos kept in *ex vivo* culture media (described earlier) with the addition of 20mM HEPES. Images were acquired with LA Vision TriM Scope II (LaVision Biotec, Germany) laser scanning microscope equipped with a Chameleon Vision II and Discovery ultrafast lasers (Coherent, USA) for different wavelengths imaged sequentially after each Z-section (Hemalatha et al., 2022). Wavelengths of 750nm were used for NAD(P)H and 940nm for H2B-GFP:Tcf-LEF or ERK-KTR reporter. Although excitation ranges of NAD(P)H and GFP overlapped, their emission was separated by band pass filters: blue range (425-475nm; NAD(P)H) and green range (500-550nm; FAD). Exclusion of nuclear GFP signal (in H2B-GFP:Tcf-LEF embryos) from NAD(P)H channel validated separation of emission signals of NAD(P)H and GFP. Embryos were imaged using a 40x water immersion lens (Nikon; N.A. 1.15) at 400 Hz with pixel size of 0.2 µm or 0.3 µm and a z-step of 1or 1.5 µm. For optimal signal to noise ratio, Line averaging of 2 was done for all NAD(P)H images. The fluorescence detected from the reduced metabolites through this method captures both NADH and NADPH (hence called NAD(P)H) (Skala et al., 2007). It should be noted that the intracellular concentrations of the non-phosphorylated NADH and NAD+ is much higher than NADPH and NADP+ (Pollak et al. 2007). We further validated that NAD(P)H fluorescence signal in the embryos closely co-localised and followed 2-NBDG uptake.

**Image analysis.** All embryos were positioned exactly along the A-P axis (refer to figures) during imaging for easier quantifications.

- ***Glut epiblast angle of expression.*** Mid-embryo sagittal sections of Glut1/3 and DAPI-stained embryos were used for glucose-uptake quantifications in Fiji/ImageJ2. For each embryo, the angle vertex was allocated at the proximal-most boundary between epiblast and extraembryonic ectoderm, at the mid-point between posterior-most (0º) and anterior-most (180º) epiblast. Two angle values were calculated for every embryo ('Glut start' and 'Glut end') to capture the range of observable Glut expression in the epiblast, along with the PS distal length of the embryo to allocate Theiler staging (ES, MS, or LS, as described above). ‘Glut start’ and ‘Glut end’ means were calculated for each stage and visualised in a rose diagram with RStudio.

- ***PS distal elongation percentage.*** Mid-embryo sagittal sections of DAPI-stained embryos were used for PS distal elongation quantifications in Fiji/ImageJ2. For each embryo, the 0% cut-off was marked by anterior epiblast morphology, the 100% cut-off was assigned to the distal-most epiblast cell, and ‘PS’ was marked at the distal-most point where the PS morphology extends (Figure 2E). Measurements were obtained for the vertical distance between 0% to 100% and 0% to ‘PS’, so that each embryo’s PS elongation percentage (0-to-PS divided by 0-to-100) is adjusted to its size. Thus, a control embryo’s PS distal elongation percentage of 95% (Figure 2E) can be interpreted as *“this embryo has elongated its PS to 95% of its epiblast height”.* Theiler stages could be assigned with this measurement: ES $\leq$ 50; 50< MS $\leq$100; LS > 100.

- ***Basement membrane (BM) breakdown.*** Mid-embryo sagittal sections of Laminin and DAPI-stained embryos were used for BM calculations in Fiji/ImageJ2. For each embryo, the 0% cut-off was marked by anterior epiblast morphology, the 100% cut-off was assigned to the distal-most epiblast cell, and ‘intact BM’ was marked at the distal-most point where laminin was still intact (Figure 3C). Measurements were obtained for the vertical distance between 0% to 100% and 0% to ‘BM’, so that each embryo’s BM breakdown percentage (0-to-BM divided by 0-to-100) is adjusted to its size. Thus, a control embryo’s BM breakdown readout of 96% (Figure 3C) can be interpreted as *“this embryo has broken down 96% of its vertical BM length”.*

- ***FITC-Fibronectin Invadopodia Assay.*** Z-sections of DAPI-stained mesoderm explants imaged with a 40X objective were used for Fibronectin perforation calculations, with $\geq$3 images captured per explant. For every image, the following measurements were calculated: cell number, and perforation number. Invadapodia degradation is represented as a percentage (perforation number divided by cell number).

- ***ERK-KTR N:C quantification.*** Sagittal sections of Erk-KTR embryos were used to quantify Erk-activity. Manual segmentations were drawn for each cell in Fiji/ImageJ2 to delineate nuclear and cytoplasmic areas, using Erk-KTR and brightfield channels to verify cell morphologies (Figure S4B). For every cell, the following measurements were quantified: nuclear area $n^a$, cytoplasmic area $c^a$ (including the nucleus), nuclear Erk-KTR intensity $n^i$, and nuclear-subtracted cytoplasmic Erk-KTR intensity $c^i$. To calculate the N:C ratio, the average nuclear intensity [$n^i/n^a$] was divided by the average cytoplasmic intensity [$c^i/(c^a-n^a)$], such that a ratio $\geq$1 indicates inactive Erk activity.

- ***Proliferation quantification.*** Live-imaged videos captured with a 40X objective were used for proliferation counts of TCF-LEF reporter mesoderm explants, quantified manually in Fiji/ImageJ2. For each explant, an overall cell count for ‘starting population’ was calculated on the first frame, and a ‘cell division’ event (TCF-LEF telophase observation) was also assigned by careful frame-to-frame assessment over the course of the video. A proliferation index (‘cell division number’ / ’starting population’) was then obtained for each explant, such that a highest index of 1 can be interpreted as “*every cell at the beginning of the video has divided by the end of the video”.* This was then adjusted to the video’s total frame length so that explants across different experimental replicates and video lengths could be compared. Thus, a control explant with a readout of 16.5% (Figure 6D) can be interpreted as "*16.5% of the mesoderm cells at the beginning of the video have divided by the end of the video*".

- ***AIVIA-based image analysis.*** AIVIA 10.5.1 AI Image Analysis Software was used to examine mesoderm explant migration dynamics. Nuclear segmentations were generated using a 'Cell Tracking' recipe, applying a pixel classifier that was trained on TCF-LEF nuclear fluorescent channels from videos of each treatment group in every experimental replicate (Figure S5B). Parameter values were modified between rounds of pixel classifier 'previewing' & training, to ensure nuclear segmentation accuracy. Every track was manually examined against the brightfield channel to verify detection accuracy. Incorrect lineages were corrected in the 'track editor' or discarded from the final dataset, and parent and daughter tracks were treated independently. Every track measurement of interest was exported to Excel and normalised to its detection length ['first frame' subtracted from 'last frame'] so that lineages of different detection lengths could be compared. Thus, all datapoints are plotted as µm/min(s).

**Quantitative RT-PCR.** Total RNA was extracted from cells using an RNeasy Micro Kit as per the manufacturer's instructions (Qiagen). cDNA synthesis was performed with 1µg of total RNA using a High-Capacity cDNA Reverse Transcription Kit according to the manufacturer's instructions (Applied Biosystems). The amounts of mRNA were measured using the PowerUp$^{TM}$ SYBR$^{TM}$ Green PCR Master Mix (Applied Biosystems). Relative levels of transcript expression were assessed by the ΔΔCt method, with *Gapdh* as an endogenous control. For qRT-PCR primers used, see Supplementary Table 2.

**Supplementary Table 2.** qRT-PCR primers used in this paper

| **Gene name** | **Forward primer** | **Reverse primer** |
|---|---|---|
| *Gapdh* | CGTATTGGGCGCCTGGTCAC | ATGATGACCCTTTTGGCTCC |
| *T/Bra* | AACTTTCCTCCATGTGCTGAGAC | TGACTTCCCAACACAAAAAGCT |
| *Eomes* | TGTTTTCGTGGAAGTGGTTCTGGC | AGGTCTGAGTCTTGGAAGGTTCATTC |
| *Mesp1* | GTCTGCAGCGGGGTGTCGTG | CGGCGGCGTCCAGGTTTCTA |
| *Pdgfra* | TGCGGGTGGACTCTGATAATGC | GTGGAACTACTGGAACCTGTCTCG |
| *Sox2* | GGCAGCTACAGCATGATGCAGGAGC | CTGGTCATGGAGTTGTACTGCAGG |
| *Pou5f1* | CTGTAGGGAGGGCTTCGGGCACTT | CTGAGGGCCAGGCAGGAGCACGAG |
| *Rex1* | CTAAAGCAAGACGAGGCAAG | AGAATGGGTTCGGAAAACTC |
| *Klf2* | CCAAGAGCTCGCACCTAAAG | GTGGCACTGAAAGGGTCTGT |
| *Gfat* | TAAGGAGATCCAGCGGTGTC | CAGCTGTCTCGCCTGATTGA |
| *Gnpnat1* | AGAAGTGGACTGGAGTCAGA | GGTCACATCTTCCACAACTG |
| *Uap1* | ACCTGCTGCAGTTCTGGAATG | CAGCTGCTCTTGATCTCTGGT |
| *Oga* | AGCGAAGATGGCAGAGGAGT | CCGTGCTCGTAAGGAAGGTA |
| *Cdh2* | CTTGAACGGAAAGTGGAATCCT | GTCAGGCTTGGAAACGTCC |
| *Mmp2* | CTCTGCGTCCTGTGCTGCCTGTTG | AAAGTGAGAATCTCCCCCAACACC |
| *Snai1* | AAGATGCACATCCGAAGCCA | CTCTTGGTGCTTGTGGAGCA |
| *Twist1* | CCGGAGACCTAGATGTCATTGT | CCACGCCCTGATTCTTGTGA |
| *Vim* | CACTAGCCGCAGCCTCTATTC | GTCCACCGAGTCTTGAAGCA |

**RNA-sequencing sample collection and data analysis.** Total RNA was extracted from mesoderm explants using a PicoPure™ RNA Isolation Kit as per the manufacturer's instructions (Applied Biosystems). RNA samples were submitted to the Yale Center for Genome Analysis for quality assessment, library preparation, and sequencing. Paired end reads were aligned using Star (v2.7.9a) to the mouse genome (GRCm38) using the Ensembl transcriptome (release 109) (Cunningham et al, 2022). Analysis of differential gene expression was performed using DESeq2 (v1.40.1). To identify differentially regulated genes between samples for downstream analyses, we selected genes with log2 fold change greater than 0.7 or less than -0.7 and an adjusted p-value <0.01. Gene ontology analysis was performed using topGO (v2.52.0) (Alexa et al, 2023). KEGG pathway analysis was performed using DAVID (https://david.ncifcrf.gov/tools.jsp) (Sherman et al., 2022; Huang et al. 2009).

**Single cell sequencing re-analysis.** Previously published single-cell RNA sequencing data from gastrulating mouse embryos were accessed using MouseGastrulationData (v1.12.0) (https://github.com/MarioniLab/MouseGastrulationData). Data were subset to only include "Epiblast," "Primitive Streak," and "Nascent mesoderm" cell states from all samples staged E6.5 to E8.5 (samples 1:10; 12:20, 23:37). Counts were log-normalized using Seurat (v. 4.3.0) (Hao et al. 2021; Stuart et al. 2019; Butler et al. 2018; Satija et al. 2015). Cells were first randomly down-sampled for visualization purposes to 400 cells per cell type. Pseudotime and trajectory interference analyses were performed using Slingshot (v2.8.0) (Street et al. 2018) for principle curve calculation; SingleCellExperiment (v1.22.0) (Amezquita et al. 2020) and scater (v.1.28.0) (McCarthy et al. 2017) were used for gene expression visualization over pseudotime. The principle curve was then traced over the first two principal components to infer pseudo-temporal organization.

**Statistics and reproducibility.** Statistical tests were performed on GraphPad Prism9.5.0 software. Two-sided unpaired t-test with Welsch's correction was applied. The same p-values were used for all the tests: ns (non-significant) $P > 0.5$, *$P < 0.05$, **$P < 0.01$, ***$P < 0.001$, ****$P < 0.0001$. Figure legends indicate the number of embryos, tissue explants, differentiation assays, and independent experiments performed for each analysis. All experiments were conducted at least two times. Statistical power calculations were not used to determine sample size.
